# Bi-steric mTORC1-selective Inhibitors activate 4EBP1 reversing MYC-induced tumorigenesis and synergize with immunotherapy

**DOI:** 10.1101/2022.02.04.478208

**Authors:** Wadie D. Mahauad-Fernandez, Yu Chi Yang, Ian Lai, Jangho Park, Lilian Yao, James W. Evans, G. Leslie Burnett, Adrian Gill, Jacqueline A. M. Smith, Mallika Singh, Dean W. Felsher

**Author notes:** Contributed equally to this work (Wadie D. Mahauad-Fernandez, Yu Chi Yang). Co-senior authors (Mallika Singh, Dean W. Felsher).

## Abstract

The MYC oncogene is causally involved in the pathogenesis of most types of human cancer but it remains therapeutically untargeted. The mTORC1 protein complex regulates cap-dependent translation through 4EBP1 and S6K and thereby, downstream MYC protein expression. However, to date, agents such as rapalogs that selectively target mTORC1 (as compared to mTORC2) fail to reactivate 4EBP1 and thus, to block MYC *in vivo.* In contrast, agents that nonselectively inhibit both protein complexes of the mTOR pathway, mTORC1 and mTORC2, can activate 4EBP1, but often suffer from a lack of tolerability including *in vivo* hepatotoxicity and immunosuppression. Here, we report the anti-tumor activity of bi-steric mTORC1-selective inhibitors, including Revolution Medicines’ clinical candidate RMC-5552, that potently and selectively target mTORC1 over mTORC2. In an autochthonous transgenic mouse model of MYC-amplified and MYC-driven hepatocellular carcinoma (HCC), representative bi-steric mTORC1-selective inhibitors suppress translation initiation via activation of 4EBP1, thereby suppressing MYC protein expression and blocking tumor growth. Furthermore, in human HCC samples, the low levels of 4EBP1 and MYC is correlated with immune reactivation. Immunohistochemistry, CIBERSORT, and CODEX reveal that selective mTORC1 inhibition results in activation of both CD4+ T cell- and NKp46+ NK cell-mediated immune surveillance. Moreover, bi-steric mTORC1-selective inhibitors synergize with α-PD-1 to induce sustained tumor regression, with immune cell degranulation and release of perforins and granzyme B. These agents also exhibit anti-tumor activity in human patient-derived xenografts of HCC, colorectal cancer, head and neck cancer, and ovarian cancer harboring genomic amplifications in *MYC*. We infer that selective mTORC1 inhibition is a potential therapeutic strategy to drive effective MYC inactivation in cancer, and the consequent restoration of immune surveillance against neoplasia.

## Introduction

The mammalian target of rapamycin (mTOR) pathway is frequently involved in the pathogenesis of human tumorigenesis including cancers of the liver, lung, kidneys, colon, prostate, and breast^1–3^. The mTOR pathway contributes to tumorigenesis by inducing cellular proliferation and survival^4^. mTOR forms two protein complexes: mTORC1 and mTORC2^1–3^. mTORC1 responds to growth and stress factors resulting in the phosphorylation of two proteins, 4E-binding protein 1 (4EBP1) which becomes inactivated and serine/threonine 6 kinase (S6K) which becomes activated. mTORC2 responds to growth factors resulting in the phosphorylation of Ak strain transforming (AKT) protein which becomes activated (Fig. 1a, left). mTORC1 induces downstream protein translation while mTORC2 induces downstream transcription and prevents protein degradation of different oncogenes including MYC (Fig. 1a, left).

**Figure 1.**
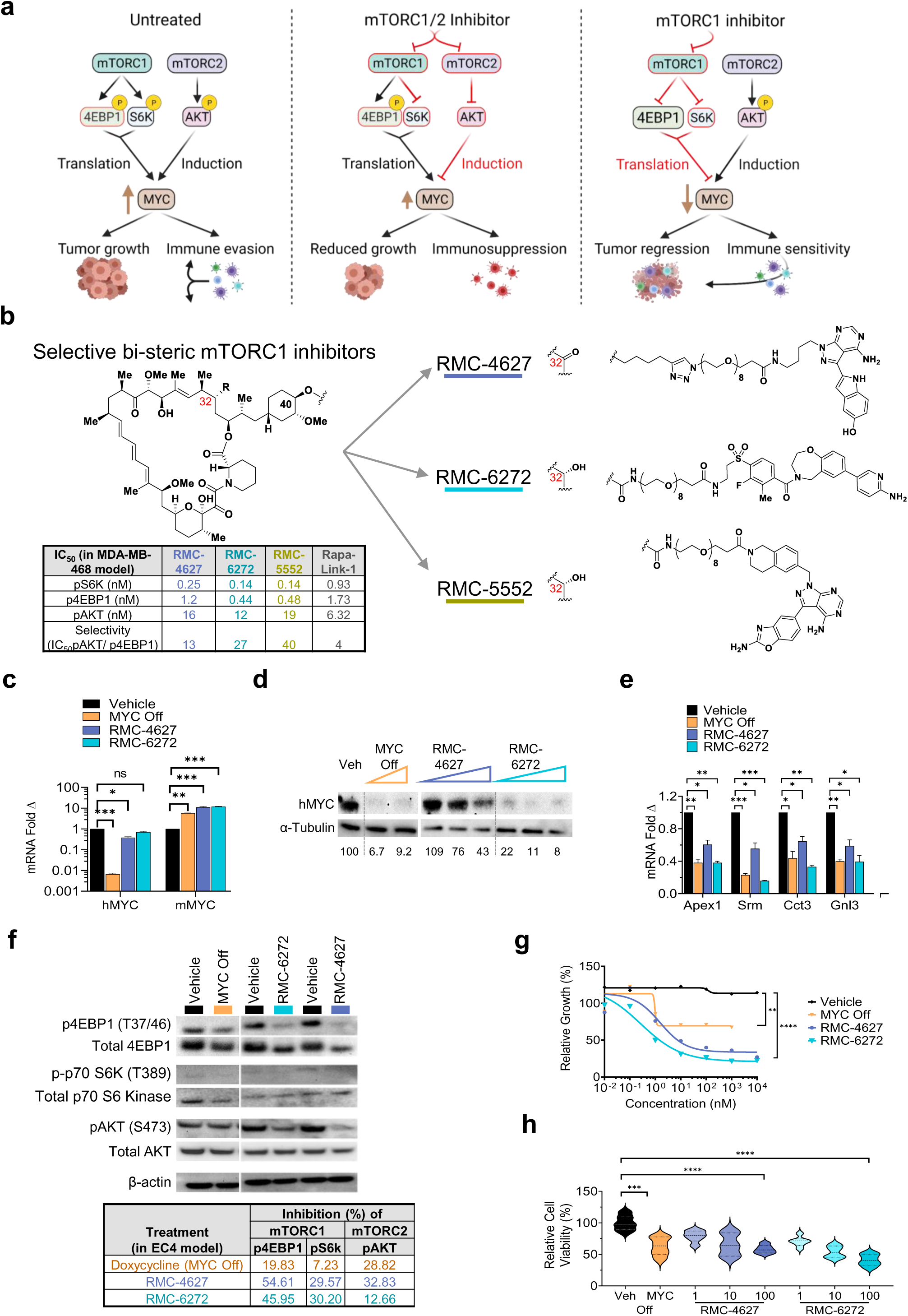
Bi-steric mTORC1-selective inhibitors reduce the growth of MYC-driven HCC cells and MYC protein levels. **a**, Graphical depiction of the study showing overactivation of mTOR signaling in cancer cells (left), the effect of current generation of mTOR inhibitors in mTORC1 and mTORC2 signaling as well as in anti-cancer immune responses (middle) and, the proposed effect of mTORC1 selective inhibitors in mTORC1 and mTORC2 signaling as well as in tumor growth and anti-cancer immunity tested in this study. **b**, Structure of the rapamycin-like core (left) which is part of the bi-steric selective mTORC1 inhibitors—RMC-4627, RMC-6272, and RMC-5552. Structures of the ATP competitive inhibitors used for each inhibitor (right). Table depicts the selectivity of each these inhibitors for mTORC1 relative to mTORC2. **c**, relative transcript levels of the human MYC transgene (hMYC) and endogenous mouse MYC (mMYC), **d**, representative western blot and relative levels of human MYC (hMYC) and alpha tubulin (α-Tubulin) protein in EC4 treated cells depicted as percent relative to vehicle, **e**, Relative mRNA levels of four MYC-regulated target genes including *Apex1*, *Srm*, *Cct3*, *and Gnl3*. **f**, Representative western blot images of mTORC1 factors p4EBP1 (T37/46), total 4EBP1, p-p70 S6K (T389), and total p70 S6 Kinase, mTORC2 factors pAKT (S473) and total AKT, and loading control beta actin (β-actin) from EC4 MYC-driven HCC cells treated with doxycycline at 20 ng/ml or with mTORC1 selective inhibitors RMC-4627 and RMC-6272 at 100 nM for 48 hours. Table in the bottom depicts the percent inhibition of pAKT, p4EBP1, and pS6 for each treatment group. **g**, Relative growth assessed by cell titer grow assay; and, **h**, Relative cell toxicity assessed by an MTT assay from EC4 cells after a 48-hour treatment with a vehicle control, doxycycline (to turn MYC Off), RMC-4627, or RMC-6272. Numbers under western blot represents relative protein levels compared to vehicle-treated cells which were set at 100%. Error bars represent SEM. Significance was taken at P<0.05 (*), P<0.01 (**), P<0.005 (***), and P<0.001 (****). ns = not significant.

Therapeutics against the mTOR pathway have been suggested as a strategy to target MYC-driven cancers^5–7^. MYC is one of the most commonly activated human oncogenes but it is yet to be directly targeted with a therapeutic agent^8^. MYC contributes to the pathogenesis of over 70% of human cancers and it is amplified in about 15% of hepatocellular carcinomas (HCC)^8, 9^. Experimentally, inhibiting MYC results in tumor regression associated with immune reactivation^8^. The mTOR pathway is also commonly activated in cancer significantly increasing MYC protein expression^1, 10–13^. Genetic or pharmacological inhibition of the mTOR pathway can reduce MYC-driven tumorigenesis in multiple mouse models^1, 10–13^. The two mTOR complexes, mTORC1 and mTORC2, together regulate MYC as well as RAS, MAPK, and PI3K/AKT^1, 12, 14^. mTORC1 regulates MYC protein translation through the phosphorylation and inactivation of 4EBP1^10, 15^ (Fig. 1a, left). However, the first generation mTORC1-selective allosteric inhibitor Rapamycin has a minimal effect on inhibiting 4EBP1 phosphorylation which then results in a modest decrease of MYC levels *in vivo*^1, 16^. At tolerated doses, the nonselective dual mTOR kinase inhibitor MLN0128 transiently reduces p4EBP1, and as a result MYC levels, causing reduced tumor growth in preclinical models^1, 16^. Altogether, existing mTOR pathway inhibitors including Rapamycin and MLN0128 fail to sufficiently activate 4EBP1 *in vivo*, and therefore cannot sufficiently decrease MYC protein levels^17^ to drive tumor regression (Fig. 1a, middle).

To date, mTOR inhibitors have limited clinical benefit in cancer patients. Everolimus, a first generation Rapamycin analog, effectively inhibits S6K phosphorylation downstream of mTORC1, but does not inhibit 4EBP1 phosphorylation, which may explain its limited therapeutic effect^18^ (Fig. 1a, middle). Second generation nonselective mTOR kinase inhibitors, such as MLN0128, can inhibit p4EBP1 but the feasible dose and clinical efficacy of these agents is limited^19^ by various toxicities, including hepatotoxicity and immunosuppression^20, 21^. The inhibition of both mTORC1 and mTORC2 causes hepatotoxicity by increasing reactive oxygen species^22^ and immunosuppression characterized by hyposplenism and reduced activation and proliferation of T and natural killer (NK) cells^18, 23–31^.

Given the importance of the mTORC1 pathway in the progression of MYC-driven cancers, we hypothesized that a potent and selective mTORC1 inhibitor that enables the activation of 4EBP1 would lead to more robust inhibition of MYC protein expression, but without hepatotoxicity or immunosuppression (Fig. 1a, right). Revolution Medicines has developed third-generation bi-steric selective inhibitors of mTORC1 with a rapamycin-like core moiety covalently linked to an mTOR active-site inhibitor^32, 33^ (Fig. 1b, left). Here, we show that these agents suppress both S6K and 4EBP1 phosphorylation, and deplete MYC protein expression *in vivo* in an autochthonous conditional transgenic mouse model of MYC-driven HCC^34–36^. Furthermore, the bi-steric mTORC1-selective inhibitors demonstrate significant anti-tumor activity as monotherapy in our MYC-driven HCC mouse model without causing hepatotoxicity or immunosuppression. Importantly, RMC-5552, a bi-steric selective mTORC1 inhibitor currently in Phase 1 (NCT04774952), exhibits anti-tumor activity in human patient-derived xenografts (PDXs) preclinical models of HCC, colorectal, head and neck, and ovarian cancer with MYC amplifications. We also show that bi-steric mTORC1-selective inhibitors restore tumor immune surveillance and synergize with α-PD-1 immune checkpoint therapy in the MYC-driven HCC mouse model. Hence, we provide proof-of-principle that selective mTORC1 pharmacological inhibition can effectively target MYC-driven cancers, combine with immune checkpoint inhibition, and induce sustained tumor regression in the preclinical setting.

## Materials and Methods

### Transgenic mice

Animals were identically raised and housed at Stanford University following Administrative Panel on Laboratory Animal Care (APLAC) guidelines and procedures. The Tet-off system transgenic lines used for conditional MYC expression and oncogene addiction have been previously described^34–36^. The Lap-tTA/Tet-O-MYC transgenic mice develop hepatocellular carcinoma within four to five weeks upon removal of doxycycline from 4 week old mice, whereby there is induced expression of the MYC transgene^36^. Mice genotypes were confirmed by PCR of genomic DNA from animal tails.

### Cell culture

The Lap-tTA/Tet-O-MYC HCC tumor-derived cell line EC4 was grown in DMEM supplemented with 10% FBS, 1% penicillin/streptomycin, 1% Glutamine, 1% non-essential amino acids, and 1% pyruvate at 37°C in 5% CO_2_. To inactivate or turn off expression of the MYC transgene, EC4 cells were treated for 48 hours with 20 ng/mL of doxycycline^37^.

### Cell viability assay

A total of 3,000 MYC-driven HCC EC4 cells were plated in clear-bottom black 96-well plates, allowed to attach overnight and treated with different concentrations of a vehicle control, RapaLink-1, RMC-4627^32^, or RMC-6272, or doxycycline for 48 hours. Upon treatment completion, EC4 cells were incubated with 5 mg/ml MTT reagent for 3.5 hours followed by addition of MTT solvent^38^ according to manufacturer’s instructions (Abcam). 96-well plates were rocked for 15 minutes and read at an absorbance at 590 nm using a SpectraMax Paradigm Multi-Mode Detection Platform plate reader (Molecular Devices).

### Cell growth assay

We plated 3,000 EC4 cells in a 96-well plates and treated with different concentrations of a vehicle control, RapaLink-1, RMC-4627, or RMC-6272, or doxycycline for 48 hours. Upon treatment completion, a CellTiTer Glow assay was performed according to manufacturer’s instructions (Promega). 96-well plates were rocked for 5 minutes at room temperature, 100 µl from each well were transferred to a white opaque 96-well plate and luminescence was read using a SpectraMax Paradigm Multi-Mode Detection Platform plate reader (Molecular Devices).

### Real-time quantitative PCR

We seeded 6-well plates with EC4 cells at 80% confluency and treated with a vehicle control, doxycycline at 20 ng/mL (MYC Off), or the inhibitors RapaLink-1, RMC-4627, or RMC-6272 at 100 nM for 48 hours at 5% CO_2_ in a 37°C incubator. EC4 cells were collected and pelleted, and RNA was isolated using the RNeasy Plus Mini Kit (Qiagen) according to manufacturer’s instructions. Equal amounts of RNA were used to synthesize cDNA using SuperScriptIII (ThermoFisher) and real-time quantitative PCR was performed using SYBR Green pPCR kit (Roche) using a HT7900 Real-Time PCR system with QuantStudio12K Flex software (Applied Biosystems). Each sample was run in triplicates and raw data was processed using the cycle threshold method were samples were normalized using UBC.

The primers used in RTqPCR analyses were: human c-MYC—forward primer 5’-CTGCGACGAGGAGGAGAACT-3’ and reverse primer 5’-GGCAGCAGCTCGAATTTCTT-3’; mouse c-MYC—forward primer 5’-TCTCCATCCTATGTTGCGGTC-3’ and reverse primer 5’- TCCAAGTAACTCGGTCATCATCT-3’; mouse UBC—forward primer 5’-ACCCAAGAACAAGCACAAGG-3’ and reverse primer 5’-AGCCCAGTGTTACCACCAAG-3’; mouse APEX1—forward primer 5’-ACGGGGAAGAACCCAAGTC-3’ and reverse primer 5’-GGTGAGGTTTTCTGATCTGGAG-3’; mouse Srm—forward primer 5’-ACATCCTCGTCTTCCGCAGTA-3’ and reverse primer 5’-GGCAGGTTGGCGATCATCT-3’; mouse Cct3—forward primer 5’-GGACCTGCTTGGGACCTAAAT-3’ and reverse primer 5’-CGGGATGCTGGACTTGAATC-3’; and, mouse GNL3—forward primer 5’-CAGGATGCTGACGATCAAGAAA-3’ and reverse primer 5’-CAGATGGCTTACCTGCTGTTG-3’.

### Western blots

One million EC4 cells were plated on 6-well plates and treated with DMSO as a vehicle control, doxycycline at 20 ng/mL (MYC Off), or the inhibitors RapaLink-1, RMC-4627, or RMC-6272 at 0, 10, or 100 nM for 48 hours. Following treatment, cells were pelleted and stored at -80°C until processed for western blots. Protein isolation, western blots, and antibodies used—human c-MYC (D84C12) #5605 and alpha tubulin (DM1A) Mouse mAb #3873 from Cell Signaling Technologies—were previously described by our group^39, 40^. Antibodies to detect p-4EBP1 (T37/46) (236B4) #2855, 4EBP1 (53H11) #9644, p-p70 S6K (T389) #9205, p70 S6 Kinase (49D7) #2708, and β-actin (8H10D10) #3700 were all from Cell Signaling Technologies. Western blot bands were quantified using ImageJ as previous described^41^.

### Size Separation “SimpleWestern” Nanoimmunoassay (NIA)^42^

Simple Western 12–230 kDa size-assays were carried out using Peggy Sue^TM^ analyzer (Protein Simple, San Jose, CA, USA). First, we prepared a 384-microwell plate containing samples mixed with fluorescent standards and denatured at 95°C during 5 min, a ladder including 6 size markers, primary antibodies (dilution 1/50 in our study), secondary antibodies, the detection solution (equivalent volumes of luminol and peroxide), stacking and separation matrices, according to the manufacturer’s preparation template. After centrifugation at 1000 xg during 5 minutes, the 384-microwell plate was loaded into Peggy Sue^TM^ platform. Then, the complete automated workflow, ranging from samples’ uptake on the plate to the chemiluminescent detection and quantification of proteins, was launched. First, samples were loaded into 12 individual single-use capillaries per cycle (1 cycle comprising 11 samples and 1 ladder), followed by a size-based separation (45 minutes in our study). A detection of fluorescence of standards, mixed with every sample, ensured a control of electrophoresis. Sample proteins were captured onto capillary beds after illumination with UV light. Incubations of primary (From Cell Signaling Technologies antibodies for c-MYC (D84C12) Rabbit mAb #5605, pS6RP (Ser235/236) (D57.2.2E) XP® Rabbit mAb #4858, p4EBP1 (Thr37/46) (236B4) Rabbit mAb #2855 and pAKT (Ser473) antibody #9271). Antibodies for beta macroglobulin [EP2978Y] (ab75853) are from Abcam) and secondary antibodies were both 30 minutes in our protocol. Finally, a chemiluminescent detection of target proteins was automatically performed with different exposure times, ranging from 4 to 512 seconds. Target proteins’ sizes were determined for each sample in comparison to the migration of the size markers of the ladder. Heights and areas of all detected peaks were systematically calculated on Compass for Simple Western® software, version 4.1.0, using the high dynamic range exposure time.

### Charge Separation Nanoimmunoassay (NIA)

NIA is a highly sensitive capillary-based isoelectric focusing method that uses antibody detection to quantify protein isoforms as well as characterize posttranslational protein modifications, such as phosphorylation. Charge separation analyses were performed according to the Protein Simple user manual. Tissue homogenate samples were lysed with Bicine/CHAPS Lysis Buffer (Protein Simple, San Jose, CA, USA) containing phosphatase and protease inhibitors. Lysates were mixed with ampholyte premix and fluorescently labeled isoelectric point (pI) standards for analysis on the Peggy Sue® system (Protein Simple) according to the manufacturer’s instructions. In brief, a mixture of 0.1 µg/µL cell lysate, 1x fluorescent pI standard ladder 1, and 1x premix 4 to 7.3 ampholyte (Protein Simple) were loaded onto the capillaries. Capillary isoelectric focusing electrophoresis was carried out at 21,000 microwatts for 40 minutes. The separated proteins were immobilized to the capillary wall by exposing to UV light for 100 seconds at instrument default setting. Immunoprobing was done using primary antibodies (ERK1/2 (Millipore), diluted 1:300 and 4EBP1 (Cell Signaling technologies) diluted 1:50, or Beta-2-microglobulin (Abcam) diluted 1:100) and then probed with horseradish peroxidase (HRP)-conjugated, goat anti-rabbit or goat anti-mouse secondary antibodies (Protein Simple). All antibody incubation and wash steps were programmed and performed automatically in the Peggy Sue® system; the incubation time was 2 hours for primary antibodies and 1 hour for secondary antibodies. A 1:1 Luminol and peroxide (Protein Simple) mixture was then added to generate chemiluminescent light, which was captured by a charge-coupled device camera. The digital image was analyzed and quantified with Compass for Simple Western® software, version 4.1.0 (Protein Simple) with peak areas corresponding with signal strength.

### *In vivo* RMC-6272 short-term treatment

Lap-tTA/Tet-O-MYC mice were taken off doxycycline to activate the human transgenic MYC oncogene 4 weeks after being born, then after waiting 4-5 weeks, mice were imaged using magnetic resonance imaging (MRI) every week to precisely define tumor volumes before treatment enrollment. Mice bearing HCC tumors with these volumes were assigned to one of the following treatment groups: vehicle control (5/5/90 Transcutol/Solutol HS 15/Water), doxycycline (MYC Off), RMC-4627, RMC-6272 or Sorafenib. For vehicle control, RMC-4627, and RMC-6272 groups, mice were given one weekly intraperitoneal injection at 10 mg/kg while mice in the Sorafenib group were treated with 30 mg/kg with a daily dose given via oral gavage. Doxycycline was supplemented in water at 100 µg/mL. Seven days after treatment, mice were given a second dose of their respective treatments and they were euthanized at 4, 12, 24, 48 or 72 hours to grossly quantify tumor nodules, weight tumors, and collect HCC tumors, plasma, and other tissues.

### *In vivo* RMC-6272 long-term treatment

Lap-tTA/Tet-O-MYC mice were taken off doxycycline to activate the human transgenic MYC oncogene 4 weeks after being born, then after waiting 4-5 weeks, mice were imaged using MRI every week to verify tumor volume before treatment enrollment. Mice bearing HCC tumors were assigned to one of the following treatment groups: vehicle control, RMC-6272, α-PD-1 (BioXCell Clone RMP1-14 and catalog number BE0146), or RMC-6272 with α-PD-1. Mice in the vehicle and RMC-6272 groups were treated with a weekly dose at 10 mg/kg injected intraperitoneally. Mice in the α-PD-1 group were intraperitoneally treated with a triweekly dose of 300 µg. Mice in the RMC-6272 + α-PD-1 group were treated with RMC-6272 at day 1 and with α-PD-1 at the doses specified above on days 2, 4, and 6. The dosing for each group was repeated weekly along with weekly MRI imaging. On day 28, mice were imaged and then given a second dose of their respective treatments and they were euthanized at 24 hours to grossly quantify tumor nodules, weight and collect HCC tumors, plasma, and other tissues.

### Magnetic Resonance Imaging (MRI)

MRI screening was utilized to enroll mice with similar tumor burden into randomized treatment groups and to track tumor progression throughout the treatments. MRI screening was performed using 7T small animal MRI (Agilent conversion) with a 40 mm Varian Millipede RF coil (ExtendMR LLC, Milpitas, CA) at the Stanford Small Animal Imaging Facility as previously described^43, 44^. Briefly, animals were anesthetized with 1-3% isoflurane and placed into the MRI scanner containing a 40 mm Varian Millipede RF coil (ExtendMR LLC, Milpitas, CA). ParaVision (PV6.01) was used to acquire the DICOM images. Once the DICOM images were acquired, tumor volumes were quantified and 3D models of tumors were constructed using Osirix image processing software (Osirix, UCLA, Los Angeles, CA). The relative area difference of the 3D models before and after treatments were quantified using FIJI ImageJ software (NIH). Percent tumor regression was calculated by dividing the number of mice whose tumors were smaller by Day 7 compared to Day 0 to the total number of mice treated in each group and multiplying this number by 100%. The same calculation was done for mice enrolled in long-term treatments. For example, regression at Day 14 was calculated by dividing the number of mice whose tumors were smaller by Day 14 compared to Day 7 to the total number of mice treated in each group and multiplying this number by 100%.

### Liver function and toxicity studies

250-300 µl of blood were collected via the retro-orbital route from Lap-tTA/Tet-O-MYC mice before treatment with RMC-4627 or RMC-6272 (Day 0). Mice were given 500 µl of saline intraperitoneally to help with blood recovery. On Day 7 after treatment with RMC-4627 or RMC-6272 at 10 mg/kg, another 250-300 µl of blood were collected from the same mice and submitted to the Animal Diagnostic Laboratory at Stanford University to measure plasma levels of different circulating factors including: alanine aminotransferase (ALT), aspartate transaminase (AST), alkaline phosphatase (Alk. Phos.), Billirubin, gamma-glutamyl transferase (GGT), cholesterol, triglycerides, sodium, potassium, high-density lipoprotein (HDL), low-density lipoprotein (LDL), chloride, CO_2_, glucose, albumin, creatinine, total protein, blood urea nitrogen (BUN), calcium, and globulin. For liver function and toxicity studies in mice treated with vehicle control, RMC-6272, α-PD-1, or RMC-6272 + α-PD-1, blood was collected retro-orbitally at day 28 post-treatment.

### Survival meta-analysis

To determine differences in progression free survival in HCC patients with high *versus* low MYC signature genes and EIF4EBP1 we downloaded The Cancer Genome Atlas (TCGA) Liver hepatocellular carcinoma (LIHC) RNA-seq dataset from xenabrowser.net. We then used the Qlucore Omics Explorer software to run the HALLMARK_MYC_TARGETS_V2 gene set (https://www.gsea-msigdb.org/gsea/msigdb/cards/HALLMARK_MYC_TARGETS_V2) and using signscore, we determined patients’ samples with high and low MYC signature. From this analysis, we extracted the top and bottom 40 samples in each group, respectively. Following separation by MYC signature, we used the top and bottom 20 samples in each group that had the highest and the lowest levels of EIF4EBP1, respectively. We then plotted their respective overall survival, disease-free survival, and progression-free survival. Similarly, to choose MYC+/EIF4EBP1+ and MYC-/EIF4EBP1-samples, we selected those with the highest and lowest EIF4EBP1 within MYC high and MYC low groups getting 14 samples per group. Within these samples per group, those samples lacking survival data were excluded from the study.

### Metanalysis of protein levels from HCC patients

To generate the volcano plot of protein levels using the TCGA LIHC and The Cancer Proteome Atlas (TCPA) LIHC datasets, we first segregated patients’ samples into high and low MYC signature using the HALLMARK_MYC_TARGETS_V2 gene set and signscore as described above. The top and bottom 20 samples were aligned to the TPGA LIHC dataset to choose the patients for which there is protein data in the TCPA LIHC dataset. There were seven patient samples in the MYC high group and 11 patient samples in the MYC low group. We used these 18 samples to calculate the -log_10_ p-value and to calculate protein levels as log_2_ MYC high over MYC low which are depicted as a volcano plot.

### CIBERSORT

We downloaded the TCGA LIHC RNA-seq dataset from xenabrowser.net. We then used the Qlucore Omics Explorer software to open this dataset and to run the HALLMARK_MYC_TARGETS_V2 gene set (https://www.gsea-msigdb.org/gsea/msigdb/cards/HALLMARK_MYC_TARGETS_V2) to segregate patients’ samples into high and low MYC signature using signscore as described above. From this analysis, we extracted the top and bottom 20 samples in each group, respectively. Following separation by MYC signature, we plugged the RNAseq data into cibersortx.stanford.edu using the LM22 gene matrix to estimate the presence of tumor infiltrating immune cell populations from HCC patient samples expressing high and low MYC signature genes. To estimate abundance of immune populations from RNA-seq data of murine HCC tumors treated with vehicle, doxycycline, RMC-4627, RMC-6272, or sorafenib, we utilized CIBERSORT^45, 46^ and TIMER2.0^47^ using the http://timer.cistrome.org/ website.

### CO-Detection by indexing (CODEX) antibody staining of Fresh Frozen mouse tissue

The tissue section was removed from -80⁰C and placed onto a bed of drierite absorbent beads. After 2 minutes the tissue was placed in acetone for 10 minutes. The tissue was then allowed to dry for 2 minutes and hydrated in hydration buffer (Akoya Biosciences) twice for 2 minutes. After hydration, tissue was fixed for 10 minutes using 1.6% paraformaldehyde (PFA) in hydration buffer. This was followed by two washes in hydration buffer and equilibration in staining buffer (Akoya Biosciences) for 20 minutes. Blocking buffer was prepared by adding N, S, J and G blocking solutions to CODEX staining buffer (Akoya Bioscience). Antibodies were then added to this blocking buffer in a 1:50 titration to make a total volume of 100 µl. The antibody cocktail was added to the coverslip and staining was performed in a sealed humidity chamber at 4⁰C overnight. After staining, coverslips were washed twice in hydration buffer for a total of 4 minutes followed by fixation in storage buffer (Akoya Bioscience) with 1.6% paraformaldehyde for 10 minutes. Coverslips were then washed thrice in PBS, followed by a 5 minute incubation in ice cold methanol on ice followed by another three 1x PBS washes. CODEX fixative solution (Akoya Bioscience) was prepared right before the final fixation step. 20 µl of CODEX fixative reagent was added to 1000 µl 1x PBS. 200 µl of the fixative solution was added to the coverslip for 20 minutes followed by 3 washes in 1x PBS. Coverslips were then stored in CODEX storage buffer at 4⁰C until imaging (for a maximum of 2 weeks). Antibodies used in the experiment are Akoya Bioscience’s validated panel for mouse fresh frozen tissue.

### CODEX multicycle setup and Imaging

Coverslips were taken out of 4⁰C and allowed to come to RT for 15-20 minutes. The coverslips were mounted onto Akoya’s custom made stage in between coverslip gaskets with the tissue side facing up. The coverslips were cleaned from the bottom using a Kim wipe to get rid of any salts. The tissue was stained with Hoechst nuclear stain at a 1:2000 dilution in 1x CODEX buffer. A 96-well plate was used to set up the multicycle experiment with different fluorescent oligonucleotides in each well. Reporter stock solution was prepared containing 1:300 Hoechst stain and 1:12 dilution of assay reagent in 1x CODEX buffer (Akoya Bioscience). Fluorescent oligonucleotides (Akoya Bioscience) were added to this reporter solution at a final concentration of 1:50 in a total of 250 µl per well. A blank cycle was performed at the start and end of the experiment. Automated image acquisition and fluidics was performed using Akoya’s software driver CODEX Instrument Manager (CIM) and the CODEX platform (Akoya Bioscience). Imaging was performed using a Keyence BZ-X800 microscope, fitted with a Nikon CFI Plan Apo 20X/0.75 objective. The navigation tool in the BZ-X software (Keyence) was used to set up the center of each of the TMA and define each region. 15 z steps were acquired with the pitch set at 1.5 in the BZ-X software. All images were processed using the CODEX processor (Akoya Bioscience). Numbers of Ki67, CD4, CD8a, CD11c, and CD19 positive cells were manually counted using imageJ.

### Patient-derived xenograft (PDX) studies

Four HCC PDXs as well as one PDX each from a colorectal cancer patient, head and neck cancer patient, and an ovarian cancer patient with MYC amplifications were grown in immunocompromised mice. Once PDXs reached a size of 200-300 mm^3^, they were treated with vehicle control or RMC-5552 at 3 or 10 mg/kg/week via intraperitoneal injection for 28 days. All PDX mouse studies were conducted in compliance with all applicable regulations and guidelines of the Institutitonal Animal Care and Use Committee (IACUC). Tumor volumes and body weights were measured and weighed, respectively, twice a week during the study.

For PDX studies conducted at GenenDesign, Shanghai, China, and at WuXi AppTec (Suzhou), China, female BALB/c nude mice were used. For PDX studies conducted at Charles River Discovery Research Services Germany GmbH, female NMRI nu/nu mice (NMRI-Foxn1nu) were used.

### RNA sequencing

#### Data Analysis

Downstream analysis was performed using a combination of programs including STAR, HTseq, Cufflink and our wrapped scripts. Alignments were parsed using the Tophat program and differential expressions were determined through DESeq2/edgeR. GO and KEGG enrichment were implemented by the ClusterProfiler. Gene fusion and difference of alternative splicing event were detected by Star-fusion and rMATS software.

#### Reads mapping to the reference genome

Reference genome and gene model annotation files were downloaded from genome website browser (NCBI/UCSC/Ensembl) directly. Indexes of the reference genome was built using STAR and paired-end clean reads were aligned to the reference genome using STAR (v2.5). STAR used the method of Maximal Mappable Prefix(MMP) which can generate a precise mapping result for junction reads.

#### Quantification of gene expression level

HTSeq v0.6.1 was used to count the read numbers mapped of each gene. And then FPKM of each gene was calculated based on the length of the gene and reads count mapped to this gene. FPKM, Reads Per Kilobase of exon model per Million mapped reads, considers the effect of sequencing depth and gene length for the reads count at the same time, and is currently the most commonly used method for estimating gene expression levels^48^.

#### Differential expression analysis

(For DESeq2 with biological replicates) Differential expression analysis between two conditions/groups (two biological replicates per condition) was performed using the DESeq2 R package (2_1.6.3). DESeq2 provide statistical routines for determining differential expression in digital gene expression data using a model based on the negative binomial distribution. The resulting P-values were adjusted using the Benjamini and Hochberg’s approach for controlling the False Discovery Rate(FDR). Genes with an adjusted P-value <0.05 found by DESeq2 were assigned as differentially expressed. (For edgeR without biological replicates) Prior to differential gene expression analysis, for each sequenced library, the read counts were adjusted by edgeR program package through one scaling normalized factor. Differential expression analysis of two conditions was performed using the edgeR R package (3.16.5). The P values were adjusted using the Benjamini & Hochberg method. Corrected P-value of 0.05 and absolute foldchange of 1 were set as the threshold for significantly differential expression. The Venn diagrams were prepared using the function vennDiagram in R based on the gene list for different group.

#### Correlations

To allow for log adjustment, genes with 0 FPKM are assigned a value of 0.001. Correlation were determined using the cor.test function in R with options set alternative =”greater” and method = “Spearman”.

#### Clustering

To identify the correlation between difference, we clustered different samples using expression level FPKM to see the correlation using hierarchical clustering distance method with the function of heatmap, SOM (Self-organization mapping) and kmeans using silhouette coefficient to adapt the optimal classification with default parameter in R. *GO and KEGG enrichment analysis of differentially expressed genes:* Gene Ontology (GO) enrichment analysis of differentially expressed genes was implemented by the clusterProfiler R package, in which gene length bias was corrected. GO terms with corrected P-value less than 0.05 were considered significantly enriched by differential expressed genes. KEGG is a database resource for understanding high-level functions and utilities of the biological system, such as the cell, the organism and the ecosystem, from molecular level information, especially large-scale molecular datasets generated by genome sequencing and other high-through put experimental technologies (http://www.genome.jp/kegg/). We used clusterProfiler R package to test the statistical enrichment of differential expression genes in KEGG pathways.

#### PPI analysis of differentially expressed genes

PPI analysis of differentially expressed genes was based on the STRING database, which contained known and predicted Protein-Protein Interactions. For the species existing in the database (like human and mouse), we constructed the networks by extracting the target gene lists from the database.

#### Differentially expressed gene annotation

TFCat and Cosmic database were used to annotate differentially expressed genes. TFCat is a curated catalog of mouse and human transcription factors (TF) based on a reliable core collection of annotations obtained by expert review of the scientific literature. COSMIC is a database designed to store and display somatic mutation information and related details which contains information relating to human cancers.

#### Data access

The high-throughput sequencing data from this study have been submitted to the NCBI Sequence Read Archive (SRA) under accession number ####.

### Immunohistochemistry (IHC)

HCC tumors from Lap-tTA/Tet-O-MYC mice treated with a vehicle control, doxycycline, RMC-4627, RMC-6272, or sorafenib for 7 days or with a vehicle control, RMC-6272, α-PD-1, or RMC-6272 + α-PD-1 for 28 days were immersed and fixed in 10% formalin for 24h and transferred to 70% ethanol. Tissues were embedded in paraffin using standard procedures on a Tissue-TEK VIP processor (Miles Scientific). From these paraffin blocks, 5 µm sections were mounted on Apex superior adhesive slides (Leica Microsystems) and stained as previously described^43^. Stained IHC sections were mounted with antifade mounting medium (Pro-Long Gold; Invitrogen) and coverslips were used to seal the slides, Images from slides were acquired at 25°C on a Zeiss Axiovert 200M inverted confocal microscope with a 40 Plan Neofluor objective using IP Lab 4.0 software (Scanalytics). Antibodies used for IHCs include: human MYC (Millipore Sigma c-Myc (EP121)), CD4 (abcam, ab183685, EPR19514), NKp46 (abcam, ab214468), NKG7 (Cell Signaling Technologies, 65507), perforin (Cell Signaling Technologies, E3W4I), granzyme B (Cell Signaling Technologies, D6E9W) and the same antibodies used in western blot and NIA for pS6RP, and p4EBP1.

### Statistical analyses

Graphpad prism (GraphPad Prism software) was used to generate graphs and to perform statistical analyses. These analyses were performed using an unpaired t-test assuming Gaussian distribution with Welch’s correction without assuming equal standard deviations for all comparisons between two different treatment groups while two-way ANOVA was used to determine significance between multiple treatment groups at different time points. For PDX studies, tumor volumes at the end of study were compared between control and treatment groups by two-way ANOVA Dunnet’s multiple comparison test. Log rank (Mantel-Cox) test was used for Kaplan-Meier survival analyses. Error bars represent standard deviations (SD) or standard error of the mean (SEM) as specified in each figure legend. A probability value (P) of 0.05 or lower was considered significant and is indicated by one asterisk (*) while P < 0.01 is indicated by two asterisks (**), P < 0.005 is indicated by three asterisks (***), and P < 0.001 is indicated by four asterisks (****).

## Results

### Bi-steric mTORC1-selective inhibitors potently reduce p4EBP1 *in vitro*

The prototype bi-steric pan-mTOR inhibitor RapaLink-1 and the early bi-steric mTORC1-selective tool compound RMC-4627 were previously reported^32, 33^. Here, we tested two additional bi-steric mTORC1-selective inhibitors: a research tool compound—RMC-6272 and Revolution Medicines’ development candidate—RMC-5552 (NCT04774952). Both exhibit improved potency and selectivity as compared to the pan-mTOR bi-steric inhibitor RapaLink-1 and to RMC-4627 ^32, 33^(Fig. 1b). RMC-4627, RMC-6272, and RMC-5552 all selectively inhibit mTORC1 (as compared to mTORC2) and consequently decreased phosphorylation of 4E-binding protein 1 (4EBP1) and serine/threonine 6 kinase (S6K) over phosphorylation of Ak strain transforming (AKT) protein in several cancer cell lines *in vitro* (Fig. 1b, table; selectivity of 13, 27, and 40 respectively).

The chemical structures of the bi-steric inhibitors described herein are shown in Fig. 1b. As previously described, RMC-4627 contains an ether chemical handle with an alkyne terminus at the C40 position on rapamycin. An azide containing PEG linker appended to a PP242-derived active-site inhibitor was then coupled to the alkyne containing chemical handle via a copper-catalyzed cycloaddition^32^. RMC-6272 and RMC-5552 exchanged this ether chemical handle for a C40 carbamate to allow for synthetic tractability, and incorporated the XL388-derived and MLN0128-derived active site inhibitors, respectively (Fig. 1b).

### Selective mTORC1 inhibition decreases MYC protein levels and inhibits tumor cell growth *in vitro*

Next, we studied the ability of these inhibitors to selectively inhibit mTORC1, decrease phosphorylation of 4EBP1 and S6K, and influence MYC protein expression in the mouse tumor-derived cell line, EC4. This cell line is derived from a Tet system conditional transgenic mouse model of human MYC-driven HCC (see Ref^36^). In this model human MYC transgene is constitutively and specifically expressed in the liver, and upon doxycycline treatment the human MYC gene is shut off (MYC Off) and human MYC protein is no longer expressed.

We evaluated the effects of bi-steric mTORC1-selective inhibitors on the expression of human *MYC* and mouse *Myc* mRNA and MYC protein expression. First and as expected, we found that for mRNA expression of human *MYC* (hMYC), RMC-4627 and RMC-6272 did not have any effect. But, exposure to each of these inhibitors was associated with a small increase in the levels of endogenous mouse *Myc* mRNA (mMYC) (Fig. 1c). This was suggestive that although *MYC* transgene mRNA expression was unchanged, these inhibitors were affecting the human MYC transgene protein expression, since MYC protein levels are inversely correlated with mRNA levels via feedback regulation of endogenous mouse *Myc* mRNA transcription^35, 36^. Indeed, when we measured MYC protein expression, we found that RMC-6272 and RMC-4627 reduced MYC levels relative to the control, consistent with the relative potency for p4EBP1 seen above (Fig. 1d, 92% and 57%, respectively). We found evidence that the decrease in MYC protein was associated with a decrease in its transcriptional activity on target genes, since RMC-6272, more than RMC-4627, reduced the expression of MYC target genes *Apex1*, *Srm*, *Cct3*, and *Gnl3*^49–53^ relative to the control (Fig. 1e, 50% and 30%, respectively).

Next, we examined the ability of the bi-steric mTORC1 inhibitors to selectively inhibit mTORC1 *versus* mTORC2 in EC4 cells, by measuring phosphorylation of 4EBP1, S6K, and AKT. Treatment with 100 nM of RMC-4627^32^ and RMC-6272 potently reduced p4EBP1 (55% and 46%, respectively) and pS6K (29.5% and 30%, respectively) but weakly decreased AKT phosphorylation (33% and 13%, respectively; Fig. 1f). These findings are consistent with the selectivity observed in MDA-MB-468 cells (Fig. 1b), which shows that RMC-6272, had a 27-fold higher selectivity for mTORC1 over mTORC2, and RMC-4627 was selective by 13-fold in MDA-MB-468 cells (Fig. 1b). Thus, RMC-6272 and RMC-4627 decreased p4EBP1, pS6K, and MYC protein levels in the EC4 murine HCC cell line.

We also examined the ability of RMC-6272 and RMC-4627, to inhibit *in vitro* tumor cell growth and viability. Both RMC-6272 and RMC-4627 effectively reduced the proliferation and viability of MYC-driven EC4 cells compared to the vehicle group (Fig. 1g-1h). Of note, RMC-6272 showed higher potency than RMC-4627 at decreasing proliferation and viability of EC4 cells (Fig.1g-1h).

### High levels of both 4EBP1 and MYC predict poor survival outcomes in human patients with HCC

Our *in vitro* results support the notion that mTORC1 and its regulation of MYC expression through 4EBP1^10, 15^ are key nodal points for HCC cell growth. Next, we performed meta-analyses of The Cancer Genome Atlas (TCGA) and The Cancer Proteome Atlas (TCPA) Liver hepatocellular carcinoma (LIHC) datasets. In human HCC, increased expression of genes in the hallmarks of MYC datatset (MYC^Sig^) was associated with high protein levels of mTORC1 factors—mTOR, RAPTOR, 4EBP1, P70S6K1, and S6, in line with findings from others^1, 11–13^ (Fig. 2a). HCC patients with tumors expressing low MYC^Sig^ showed better overall survival, progression-free survival, and disease-free survival than patients with high MYC^Sig^, and high *EIF4EBP1* levels further drove poorer survival in the low MYC^Sig^ patients (Fig. 2b and Sup. Fig. 1a-1b). The combination of low MYC^Sig^ and low *EIF4EBP* levels in human HCC tumors is associated with the best survival in HCC patients from our meta-analysis based on MYC^sig^ and *EIF4EBP1* expression levels.

**Figure 2.**
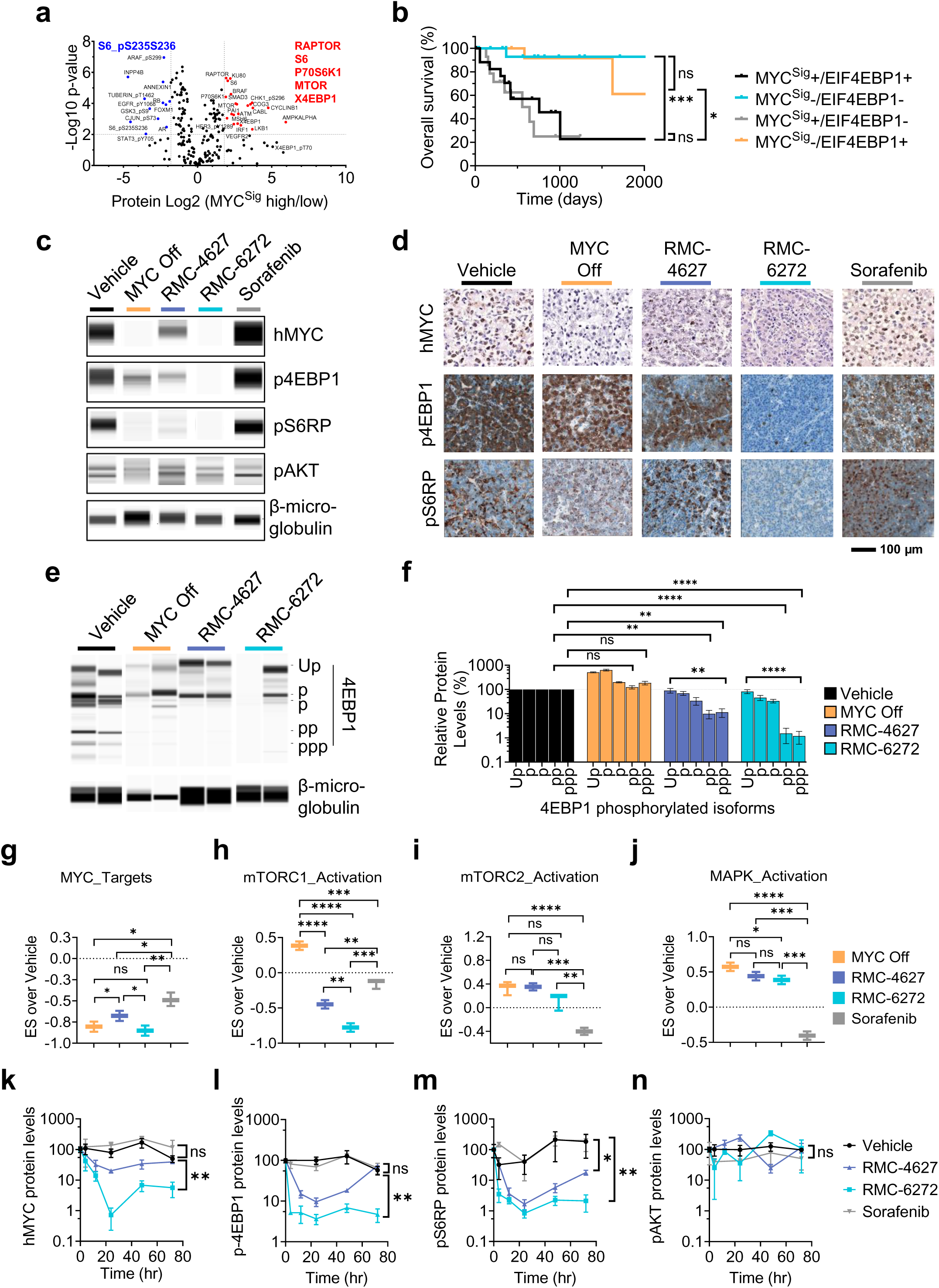
RMC-6272 has a long-lasting inhibitory effect on tumor p4EBP1 levels and MYC protein levels *in vivo*. **a**, Volcano plot of proteins that are significantly overexpressed (red) or underexpressed (blue) in MYC signature (MYC^Sig^) high *versus* low human HCC tumors (TCPA LIHC). Names of proteins which are specific to the mTORC1 pathway are highlighted in red. **b**, Overall survival of human HCC patients (TCGA LIHC) after stratifying them into those expressing MYC^Sig^ high (MYC^Sig^+) and low (MYC^Sig^-) and expressing high or low levels of *EIF4EBP1* (n=20 for MYC ^Sig^+/4EBP1+ and MYC ^Sig^-/4EBP1-; and, n=14 for MYC ^Sig^+/4EBP1- and MYC ^Sig^-/4EBP1+). **c**, Representative western blot-like images generated from size separation nanoimmunoassay (NIA), **d**, representative immunohistochemistry (IHC) images, **e**, representative NIA western blot-like images, and **f**, quantification of the different 4EBP1 phosphorylation isoforms from charge separation NIA. **g-j**, Enrichment scores (ES) of **g**, hallmarks of MYC targets geneset, **h**, hallmarks of mTORC1 signaling geneset, **i**, hallmarks of PI3K, AKT, mTOR signaling geneset, and **j**, activated MAPK activity geneset; and **k-n**, quantification of **k**, hMYC, **l**, p4EBP1, **m**, pS6RP, **n**, pAKT over β-macroglobulin protein from NIA analysis, from HCC tumors of Lap-tTA/Tet-O-MYC mice treated for 7 days with a vehicle control (5/5/90 Transcutol/Solutol HS 15/Water) injected intraperitoneally once weekly, doxycycline at 100 ug/ml in water (MYC Off), sorafenib at 30 mg/kg given via oral gavage once daily for a week or two bi-steric mTORC1-selective inhibitors RMC-4627 and RMC-6272 at 10 mg/kg intraperitoneally once weekly. At day 7, mice were treated with one more dose of their respective treatment and euthanized at 4, 12, 24, 48, or 72 hours after. In panels **c** and **d**, the number of “p”s presiding the protein name corresponds to the relative phosphorylation levels. Thus, ppp-4EBP1 is more phosphorylated than the p-4EBP1 isoform; Up = unphosphorylated. Error bars represent SEM. Significance was taken at P<0.05 (*), P<0.01 (**), P<0.005 (***), P<0.001 (****). ns = not significant. Scale bar in panel **d** is 100 µm.

### Selective mTORC1 inhibition depletes p4EBP1 and MYC protein levels *in vivo*

We evaluated the ability of RMC-4627 and RMC-6272 to activate 4EBP1 and reduce MYC protein *in vivo*. Transgenic mice with MYC-driven HCC were treated with RMC-4627 or RMC-6272, each at a single dose of 10 mg/kg/week. As assessed by nanoimmmuno assay (NIA^42^) and immunohistochemistry (IHC), RMC-6272 treatment depleted MYC protein in tumors similarly to the MYC Off control group *in vivo* while RMC-4627 reduced MYC levels by 50% as assessed using both assays (Fig. 2c-2d and Sup. Fig. 2a-2b). Also, RMC-6272 depleted p4EBP1 and pS6RP (as a surrogate marker for pS6K), but not pAKT, in MYC-driven tumors while RMC-4627 reduced p4EBP1 and pS6RP proteins by 60% and 80%, respectively, as assessed by NIA and IHC (Fig. 2c-2d and Sup. Fig. 2a-2b). Thus, RMC-4627 and RMC-6272 retain selectivity for mTORC1 over mTORC2 *in vivo*. To benchmark these results, we also examined the effect of the pan-RTK inhibitor sorafenib^54^, currently approved for HCC treatment, on reducing protein levels of MYC and mTOR substrate phosphorylation. Sorafenib treatment did not reduce MYC, p4EBP1 or pS6RP levels *in vivo* but decreased pAKT levels by 50% (Fig. 2c-2d and Sup. Fig. 2a).

Furthermore, using charge separation based NIA^42^, RMC-4627 and RMC-6272 reduced the levels of multiple heavily phosphorylated 4EBP1 isoforms without affecting levels of unphosphorylated 4EBP1 (Up-4EBP1) (Fig. 2e-2f). Notably, neither RMC-4627 nor RMC-6272 affected phosphorylation of ERK1/2, which are not regulated by mTORC1 (Sup. Fig. 2c-2d). Thus, RMC-6272, and (to a lesser extent) RMC-4627, selectively reduced p4EBP1 protein levels *in vivo* resulting in a decrease in MYC protein.

### mTORC1 inhibition potently reduces MYC transcriptional activity *in vivo*

We next examined the effect of RMC-4627 and RMC-6272 on downstream MYC transcriptional activity *in vivo.* Consistent with the effects on MYC protein levels above, RMC-6272 reduced the expression of MYC target genes and led to a relative enrichment score (ES) of -0.85, which is similar to MYC Off, while RMC-4627 also reduced their expression but to a lesser extent than RMC-6272 (Fig. 2g and Sup. Fig. 3). MYC-regulated genes can also be regulated by other oncogenes (WNT, RAS, HRAS, KRAS, BRAF, and PI3K) overexpressed in HCC^4^. To determine the specificity of RMC-6272 and RMC-4627 for MYC, we asked if these treatments decreased the expression of genes regulated by other oncogenes in HCC, via GSEA. RMC-6272, and similarly RMC-4627, did not significantly downregulate the expression of genes regulated by WNT, NRAS, HRAS, KRAS, BRAF, and PI3K (Sup. Fig. 4). Thus, RMC-4627 and to a greater extent RMC-6272 selectively reduced MYC transcriptional activity *in vivo* without influencing the transcriptional activity of other oncogenes known to be overexpressed in HCC. These results are consistent with the notion that MYC as an oncogene relying on cap-dependent translation is susceptible to inhibition of p4EBP1^15^, which was achieved by RMC-4627 and RMC-6272.

Next, we evaluated whether the selectivity of RMC-4627 and RMC-6272 for mTORC1 *versus* mTORC2 resulted in selective regulation of downstream genes using ES and GSEA *in vivo*. RMC-6272, and to a lesser extent, RMC-4627 reduced the expression of genes commonly overexpressed upon mTORC1 activation without affecting expression of genes upregulated by mTORC2 activation (Fig. 2h-2i and Sup. Fig. 3). Notably, the tyrosine kinase inhibitor approved for treatment of HCC, sorafenib^54^, decreased expression of mTORC2-regulated genes without affecting the expression of genes upregulated by mTORC1 activation (Fig. 2h-2i and Sup. Fig. 3), As expected, sorafenib but not RMC-6272 or RMC-4627, decreased the expression of MAPK-regulated genes (Fig. 2j and Sup. Fig. 3). Thus, bi-steric mTORC1-selective inhibitors, but not sorafenib, selectively reduced mTORC1 activation without affecting mTORC2 activation *in vivo*.

### Bi-steric mTORC1 inhibitor treatment results in sustained inhibition of p4EBP1 and MYC *in vivo*

We evaluated the *in vivo* durability of response for RMC-4627 and RMC-6272 to reduce p4EBP1 and MYC levels following administration of a single dose. The kinetics of MYC, p4EBP1, pS6RP, and pAKT protein inhibition by mTORC1 inhibitors was measured using NIA and IHC. As assessed by NIA, RMC-6272 rapidly reduced p4EBP1 and pS6RP by 4 hours followed by a significant reduction in MYC protein levels by 12 hours reaching its peak of inhibition by 24 hours. At this time point, RMC-6272 decreased MYC levels to 2%, p4EBP1 levels to 1% and pS6 levels to 1%. Levels of MYC, p4EBP1, and pS6RP increased slightly after 24 hours but remained relatively low (6%, 5%, and 2%, respectively) out to 72 hours (Fig. 2k-2m). RMC-4627 reduced p4EBP1, pS6RP, and MYC levels by 12 hours and reach its peak of inhibition by 24 hours decreasing MYC to 25%, p4EBP1 to 10% and pS6 to 2%. By 72 hours post RMC-4627 treatment, p4EBP1 and MYC levels recovered to 70% and 55%, respectively (Fig. 2k-2m). These results were corroborated using immunohistochemistry (Sup. Fig. 5). Importantly, pAKT levels were not reduced over a 72-hour period following treatment with either RMC-6272 or RMC-4627 (Fig. 2n). In contrast, sorafenib treatment did not reduce protein levels of MYC, p4EBP1, or pS6RP as assessed by NIA and IHC (Fig. 2k-2n and Sup. Fig. 5). Our findings show that bi-steric mTORC1-selective inhibitors exhibit potent, selective, and durable *in vivo* suppression on p4EBP1 and MYC.

### Selective mTORC1 inhibitors induce tumor regression in *MYC*-amplified human PDX models

In order to translate our findings with RMC-4627 and RMC-6272, RMC-5552 (clinical development candidate currently in Phase 1, NCT04774952) was tested in four patient-derived xenograft (PDX) models of human HCC with different *MYC* copy number variations (CNV) ranging from 2 (normal) to 6.42 (Fig. 3a-3d). RMC-5552 at two different doses did not significantly reduce tumor growth of an HCC tumor with normal *MYC* copy number (CNV = 2) (Fig. 3a). In contrast, human HCC PDX tumors with genomic amplification of *MYC* indicated by high CNV were sensitive to RMC-5552 which reduced tumor growth at tested doses (Fig. 3b-3d). Furthermore, we tested whether we could extend the proof of concept that selective mTORC1 inhibition via RMC-5552 observed in MYC-driven HCC models could reduce tumor growth in additional models of MYC-driven human cancers. Consistent with our findings in multiple HCC models, RMC-5552 was effective at reducing tumor growth of multiple human cancer models with genomic *MYC* amplifications including colorectal cancer (CRC), head and neck squamous cell cancer, and ovarian cancer in a dose-dependent manner (Sup. Fig. 6). Thus, the selective mTORC1 inhibitor RMC-5552 exhibited anti-tumor activity across a variety of human PDX models with genomic *MYC* amplification.

**Figure 3.**
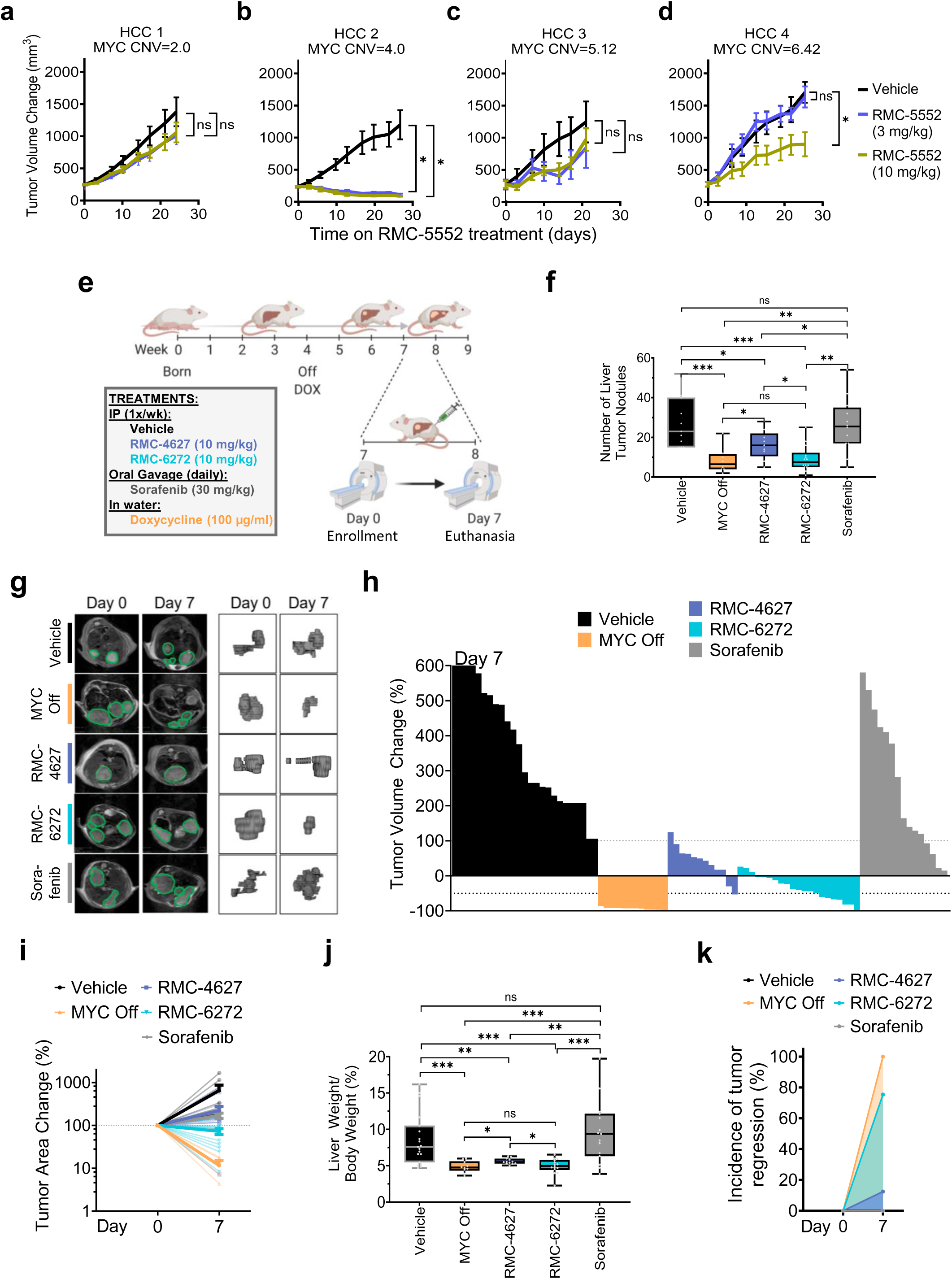
Bi-steric mTORC1 inhibitors induce regression of human and mouse MYC-driven HCC tumors. **a-d**, Tumor volume over time of four HCC patient-derived xenografts (PDXs) with different copy number variants (CNV) of MYC ranging from low (left) to high (right) treated with a vehicle control (5/5/90 Transcutol/Solutol HS 15/Water) or RMC-5552 given intraperitoneally at 3 or 10 mg/kg/week. **e**, Graphical depiction of the methods. Lap-tTA/Tet-O-MYC mice were taken off doxycycline (DOX) to induce hMYC expression and induce hepatocellular carcinoma initiation. Three weeks after, mice were MRI imaged to assess tumor volumes before enrolling them in different treatment groups which include vehicle control (5/5/90 Transcutol/Solutol HS 15/Water) injected intraperitoneally once weekly, doxycycline given at 100 µg/ml in water (MYC Off), sorafenib at 30 mg/kg given via daily oral gavage and two selective mTORC1 inhibitors RMC-4627 and RMC-6272 at 10 mg/kg weekly. **f**, Number of liver tumor nodules, **g**, Representative MRI images (left) and 3D HCC tumors (right), **h**, tumor volume change relative to Day 0 for each treatment group, **i**, tumor area change over time depicted as a percent and calculated from 3D HCC tumors before (Day 0) and after (Day 7) treatment, **j**, liver weight over body weight, and **k**, incidence of tumor regression of Lap-tTA/Tet-O-MYC mice treated with a vehicle control, doxycycline, sorafenib, RMC-4627, or RMC-6272 for 7 days. Error bars represent minimum and maximum whiskers. Tumor volumes at the end of study was compared between control and treatment groups by two-way ANOVA Dunnet’s multiple comparison test. Significance was taken at P<0.05 (*), P<0.01 (**), and P<0.005 (***). ns = not significant.

### Bi-steric mTORC1 selective inhibitors induce regression of autochthonous MYC-driven HCC tumors *in vivo*

Given the significant ani-tumor activity of RMC-5552 and the profound effects of RMC-4627 and RMC-6272 treatment on mTORC1 inhibition and consequent effects on MYC; we examined the anti-tumor activity of these bi-steric mTORC1 inhibitors following a single dose in our MYC-driven HCC conditional transgenic mouse model^36^, as compared to sorafenib dosed daily for one week (Fig. 3e). Consistent with on-target pathway suppression, as well as with MYC protein reduction described above, RMC-4627 and RMC-6272 treatment reduced the number of liver tumor nodules by 28% and 68% respectively, compared to the vehicle control, while sorafenib had no effect (Fig. 3f). RMC-6272, similar to the MYC Off group, and RMC-4627 to a lesser extent, induced regression of MYC-driven HCC tumors while the control vehicle and sorafenib did not, as assessed by tumor volume change over time calculated from 2D MRI images, and tumor area change over time calculated from 3D reconstructions of MRI images (Fig. 3g-3i and Sup. Table 1). RMC-4627 and RMC-6272 both reduced total tumor burden over a period of one week as measured by percent liver weight/body weight whereas sorafenib had no effect (Fig. 3j).

Furthermore, a single dose of RMC-6272 induced tumor regressions in 80% of treated mice while RMC-4627 induced regessions in 20% of mice compared to sorafenib which did not induce tumor regression at all (Fig. 3k). As expected, MYC Off resulted in 100% tumor regression in all animals (Fig. 3k). Impressively, the bi-steric mTORC1 inhibitor RMC-6272 induced tumor regressions in transgenic mice with large tumor burden (Sup. Fig. 7). Therefore, bi-steric mTORC1-selective inhibitor treatment showed remarkable anti-tumor activity and drove significant tumor regression in a MYC-driven HCC model following a single dose.

### Selectively inhibiting mTORC1 is well tolerated and does not induce hepatotoxicity or hyposplenism in mice harboring MYC-driven HCC

In preclinical and clinical experiments, treatment with nonselective mTOR inhibitors was associated with body weight loss, hepatic toxicity^22^, and immunosuppression characterized by hyposplenism^31^. In contrast, a single dose of RMC-6272 or RMC-4627 did not result in body weight reduction while exhibiting anti-tumor activity in a MYC-driven HCC model whereas sorafenib reduced body weight by 8% (Sup. Fig. 8a). Importantly, RMC-6272 and RMC-4627 did not induce hepatoxicity as measured through normal circulating levels of liver enzymes: ALT, AST, alkaline phosphatase, bilirubin, and GGT (Sup. Fig. 8b). These mTORC1 inhibitors did induce a modest decrease in circulating triglycerides and low-density lipoprotein (LDL) (Sup. Fig. 8c-8q). Neither RMC-6272 nor RMC-4627 were associated with hyposplenism while sorafenib reduced spleen sizes by 32% (Sup. Fig. 9). Thus, single dose administration of RMC-6272 and RMC-4627 was not associated with weight loss, hepatotoxicity or hyposplenism while driving marked anti-tumor activity in a MYC-driven tumor model.

### Downregulation of MYC signaling and *EIF4EBP1* in human HCC is predictive of immune reactivation

First-generation non-selective mTOR inhibitors are associated with immune suppression^22, 31^. But, since MYC inactivation causes immune reactivation^8^, we first evaluated if MYC signaling (MYC^Sig^) and *EIF4EBP1* levels predict immune activation in human HCC tumors using CIBERSORT^45^. Human HCC tumors expressing low MYC^Sig^ and low *EIF4EBP1 versus* those expressing high MYC^Sig^ and high *EIF4EBP1* levels had lower numbers of inactivated M0 macrophages and immune inhibitory T_reg_ cells and increased activated DCs, CD4+ memory T cells, and activated NK cells (Sup. Fig. 10). Thus, low MYC^Sig^ and low *EIF4EBP1* levels are associated with immune activation in human HCC through the recruitment of DCs, T cells, and NK cells. Based on these data, we subsequently evaluated whether RMC-6272 treatment can restore anti-tumor immunity in MYC-driven HCC tumors.

### Selective mTORC1 inhibition induces immune activation in MYC-driven tumors

We focused our continued studies on RMC-6272 because it durably suppressed MYC protein levels *in vivo*. We examined how RMC-6272 treatment influenced the immune state *in vivo* in our MYC-driven HCC mouse model through three methods: GSEA and GO Term enrichment analyses of RNA-seq data, CIBERSORT^46^ and TIMER2.0^47^, and CO-Detection by indexing (CODEX)^55^. We compared RMC-6272 with sorafenib and the MYC Off condition in select analyses as warranted.

By GSEA, RMC-6272, similarly to MYC Off, upregulated genes involved in immune system expansion and function by 24 hours which was durable through 72 hours (Fig. 4a-4c). Conversely, sorafenib did not induce genes involved in immune system expansion at any time point tested (Fig. 4a-4c). By GO Term enrichment analyses, RMC-6272 induced pathways that regulate DNA replication, cell cycle progression, and ribosome biosynthesis by 24 hours (Fig. 4d) and by 48 and 72 hours, enrichment in inflammatory responses including leukocyte migration and activation (Fig. 4e and 4f). Thus, RMC-6272 treatment may induce immune cell migration, activation, and expansion.

**Figure 4.**
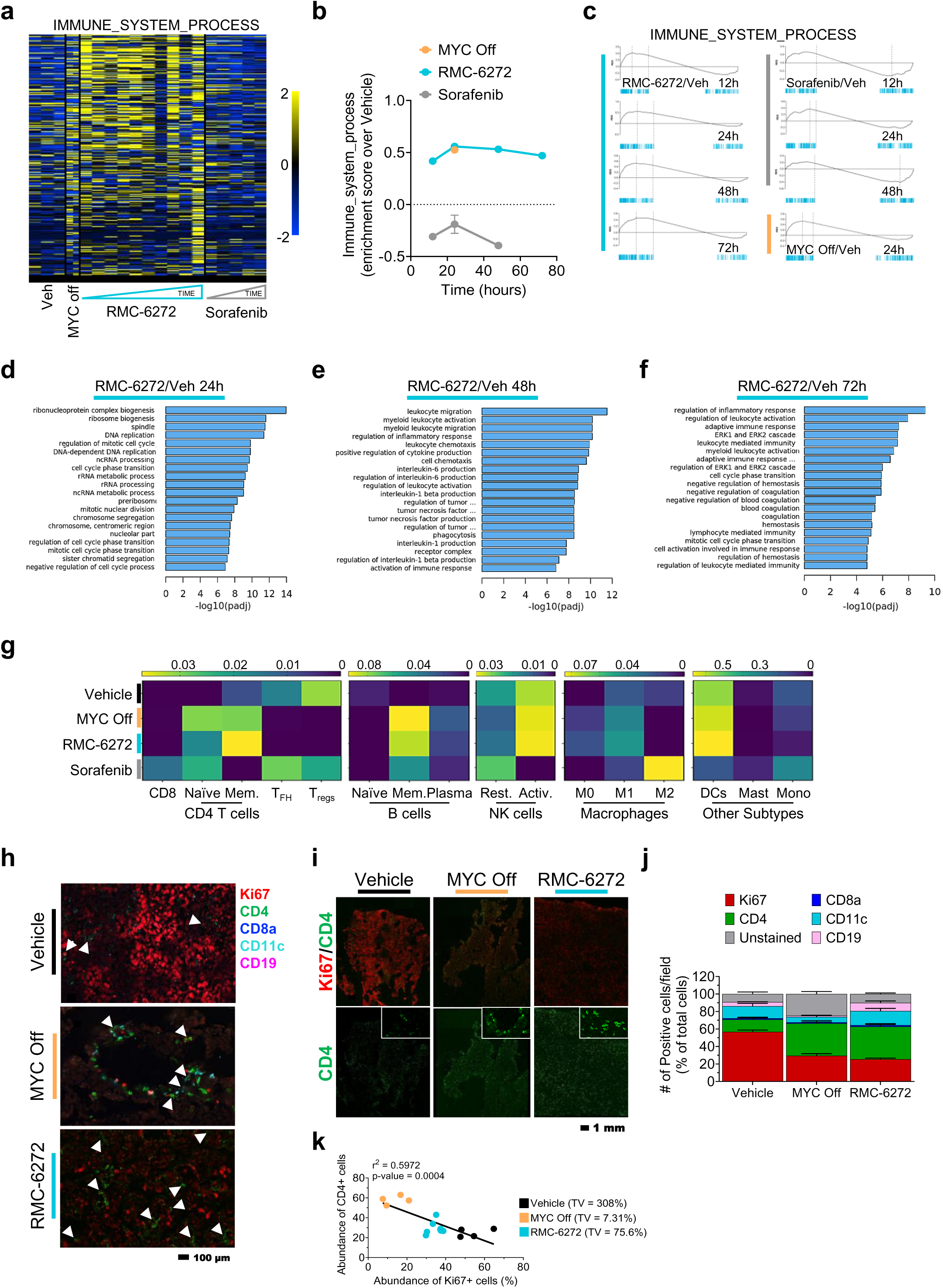
RMC-6272 induces immune activation in MYC-driven HCC tumors. **a**, Heat map of genes in the Immune System Process geneset, **b**, Enrichment scores over time from the Immune System Process geneset, **c**, GSEA plots from the Immune System Process geneset over time, and **d-f**, GO terms enrichment analysis from transcriptome data of HCC tumors of Lap-tTA/Tet-O-MYC mice treated with a vehicle control (5/5/90 Transcutol/Solutol HS 15/Water) injected intraperitoneally once weekly, doxycycline at 100 ug/ml in water (MYC Off), sorafenib at 30 mg/kg given via daily oral gavage, or the bi-steric mTORC1-selective inhibitor RMC-6272 injected intraperitoneally at 10 mg/kg once weekly. Mice were euthanized for sample collections at the following time points after a final dose on Day 7, whereas for the bi-steric inhibitors, a second dose was given followed by euthanasia and sample collections at **d**, 24, **e**, 48, and **f**, 72 hours post-dosing. **g**, Heat map of the relative abundance of different immune populations in HCC tumors of mice treated with vehicle, doxycycline (MYC Off), RMC-4627, RMC-6272, or sorafenib using CIBERSORT and TIMER2.0. The sum of the abundances of all the different immune subsets per sample equals one. **h** and **i**, Representative **h**, high magnification, and **i**, low magnification images from CODEX analysis, and **j**, quantification of protein levels of 5 markers (Ki67, CD4, CD8a, CD11c, and CD19) in HCC tumors from Lap-tTA/Tet-O-MYC mice treated for 7 days with a vehicle control, doxycycline (MYC Off), or RMC-6272. White arrow heads, in panel **h**, depict CD4+ immune cells that are present in areas with Ki67^low/-^ cells. **k**, Correlation analyses of abundance of CD4+ cells relative to Ki67+ cells quantified from CODEX data from HCC tumors of Lap-tTA/Tet-O-MYC mice treated with a vehicle control (black), RMC-6272 (light blue) or doxycycline (MYC Off, orange). Numbers next to legend correspond to the tumor volume change at Day 7 post treatment depicted as a percent relative to Day 0. Error bars represent SEM. Scale bar in panel **h** is 100 µm and in panel **i** it si 1 mm.

Second, by CIBERSORT^46^ and TIMER2.0^47^ we found RMC-6272, similar to MYC Off, decreased immune-suppressive subsets including T_reg_ cells, M2 macrophages, and mast cells in our MYC-driven HCC model (Fig. 4g). Furthermore, RMC-6272 increased memory B cells, CD4+ T cells, activated NK cells, M1 macrophages, and myeloid dendritic cells (DCs) (Fig. 4g). Moreover, treatment with sorafenib reduced memory CD4+ T cells, activated NK cells, and myeloid DCs while memory B cells increased but not to the extent of RMC-6272 or MYC Off (Fig. 4g). Notably, the abundance of T_reg_ cells and anti-inflammatory M2 macrophages increased upon sorafenib treatment in clear contrast to RMC-6272 treatment (Fig. 4g).

Third, by CODEX^55^, RMC-6272 treatment resulted in the recruitment of CD4+ cells and CD11c+ immune cells accompanied by a decrease in Ki67+ cells (Fig. 4h-4j). In RMC-6272 and MYC Off tumors, CD4+ immune cells were found in pockets of space that are cleared of Ki67+ cells or that have cells expressing low Ki67 (Fig. 4h, white arrows). Higher numbers of tumor-recruited CD4+ immune cells can be found in tissue pockets with lower numbers of Ki67+ cells; in fact, the number of CD4+ cells are inversely correlated to the number of Ki67+ cells in MYC-driven HCC tumors (Fig. 4k). Thus, RMC-6272 induced immune activation associated with both increased CD4+ and CD11c+ immune cells at sites of nonproliferating Ki67^low/-^ cancer cells.

### RMC-6272 and α-PD-1 cooperate to induce sustained tumor regression

Since RMC-6272 enhanced immune activation, we examined whether this agent could cooperate with immune checkpoint therapy. Animals with MYC-driven HCC tumors were staged and treated with a weekly dose of RMC-6272 followed by triweekly doses of α-PD-1 for 28 days (Fig. 5a). Of note, tumors in our autochthonous MYC-driven HCC mouse model do not respond to α-PD-1 therapy as shown in Fig. 5b-5c, similar to clinical reports using single immune checkpoint inhibitors in HCC^56–58^. Treatment with RMC-6272 alone or in combination with α-PD-1 markedly reduced the number of tumor nodules and the percent liver weight/body weight compared to controls or single agent α-PD-1 treatment (Fig. 5b-5c). RMC-6272 treatment effects at Day 7 were consistent with those observed in the single dose study (Fig. 3f-3k and Fig. 5d-5f). Interestingly, in the 28-day efficacy study, we observed on-treatment tumor volume increase in theRMC-6272-treated mice, with 8 of 11 mice showing tumor volume increase from baseline over time as assessed by tumor volume change over time calculated from 2D MRI images, and tumor area change over time calculated from 3D reconstructions of MRI images (Fig. 5e and Sup. Fig. 11). In contrast, only 1 of 11 (9%) mice treated with the combination therapy showed a slight tumor volume increase by week 4, indicating a combinatorial impact on treatment durability. Another indicator of a clear combinatorial effect of RMC-6272 with α-PD-1 was the increased tumor regression compared to either RMC-6272 or α-PD-1 alone (38%, 18%, and 0%, respectively, Fig. 5f, and Sup. Table 2). Furthermore, using tumor volume doubling as a surrogate for disease progression, combination of RMC-6272 with α-PD-1 significantly improved progression-free survival compared to either agent alone (Fig. 5g). Long-term treatment with RMC-6272 alone or in combination with α-PD-1 was not associated with body weight loss or hepatotoxicity (Sup. Fig. 12). Thus, combining a selective bi-steric mTORC1 inhibitor with α-PD-1 treatment induced sustained tumor regressions in a MYC-driven HCC model.

**Figure 5.**
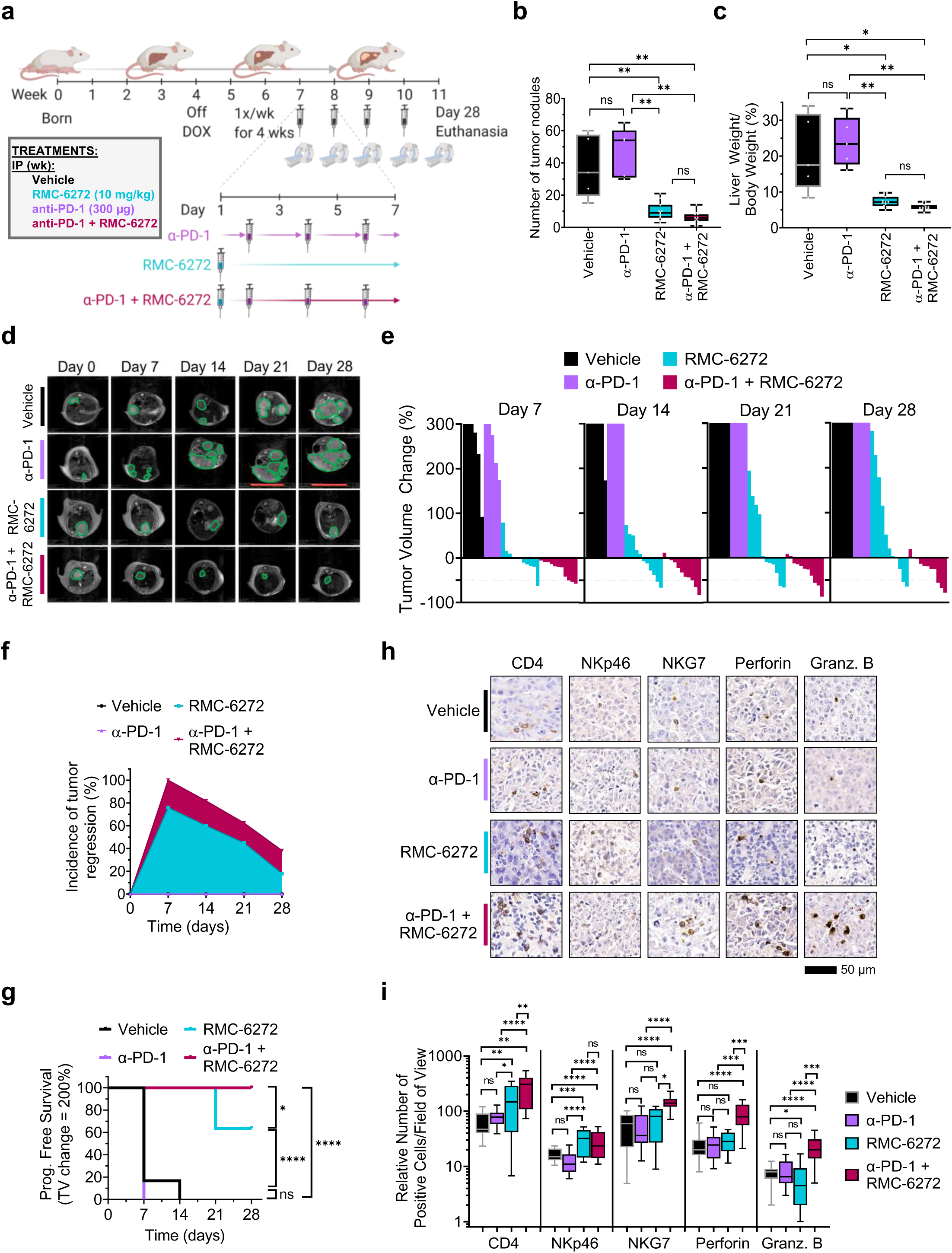
RMC-6272 in combination with α-PD-1 induces immune degranulation and sustained regression of MYC-driven tumors. **a**, Graphical depiction of the methods. Lap-tTA/Tet-O-MYC mice were taken off doxycycline (DOX) to induce hMYC expression and induce hepatocellular carcinoma initiation. Three weeks after, mice were MRI imaged to assess tumor volumes before enrolling them to be treated with vehicle (5/5/90 Transcutol/Solutol HS 15/Water) injected intraperitoneally once weekly, RMC-6272 given intraperitoneally at 10 mg/kg/week, α-PD-1 given intraperitoneally at 300 µg triweekly, or a combination of RMC-6272 with α-PD-1. **b**, Number of liver tumor nodules at day 28, **c**, liver weight over body weight at day 28, **d**, representative MRI images over time, **e**, tumor volume change over time relative to Day 0 for each treatment group, **f**, incidence of tumor regression over time, **g**, progression-free survival, and **h**, representative immunohistochemistry images, and **i**, quantification of CD4, NKp46, NKG7, perforins, and granzyme B (Granz. B) in HCC tumors from Lap-tTA/Tet-O-MYC treated for 28 days with a vehicle control (5/5/90 Transcutol/Solutol HS 15/Water), RMC-6272 at 10 mg/kg injected intraperitoneally once weekly, α-PD-1 injected intraperitoneally at 300 ug triweekly, or a combination of RMC-6272 and α-PD-1. Error bars represent minimum and maximum whiskers. The numbers in panel **i** represent the relative number of positive cells per field of view for different areas of tumors from mice in each treatment group. Log rank (Mantel-Cox) test was used for Kaplan-Meier survival analyses to compare indicated groups. Significance was taken at P<0.05 (*), P<0.01 (**), and P<0.001 (****). ns = not significant. Scale bar in panel **h** is 50 µm.

### The combination of RMC-6272 with α-PD-1 increased T and NK cell recruitment and the release of perforins and granzymes

We next examined potential mechanisms of the combinatorial effects on tumor suppression described above. Immune checkpoint therapy is associated with immune activation including T cell and NK cell degranulation and the release of perforins and granzymes to control tumor growth^59, 60^. We found that single agent RMC-6272 and more so, RMC-6272 and α-PD-1, increase intratumoral abundance of CD4+ and NKp46+ (NK cell marker) immune cells in MYC-driven tumors compared to vehicle- or α-PD-1-treated tumors suggesting T and NK cell activation (Fig. 5h-5i). RMC-6272 with α-PD-1 increased NKG7 levels compared to α-PD-1 or RMC-6272 alone even though RMC-6272 alone enhanced recruitment of CD4+ and NKp46+ cells, consistent with increased degranulation (Fig. 5h-5i). RMC-6272 with α-PD-1, more than either agent alone, increased levels of perforins and granzyme B (Fig. 5h-5i). Thus, RMC-6272 with α-PD-1 increased recruitment of CD4+ cells, immune cell degranulation, and release of perforins and granzyme B in MYC-driven tumors, relative to single agent RMC-6272, and consistent with the enhanced anti-tumor effects resulting from the combination therapy.

## Discussion

The mTOR pathway is key to regulating the protein translation of MYC. MYC is global a driver of cancer cell growth and immune evasion, but remains to be therapeutically targeted^8^. The inhibition of the mTOR pathway, in particular mTORC1 and leading to downstream activation of 4EBP1, and the consequent inhibition of cap-dependent translation, has therapeutic potential in MYC-driven tumors^15^. First generation mTORC1 selective inhibitors such as Rapalogs are unable to reactivate 4EBP1, have shown limited clinical efficacy, and also appear to have dose limiting hepatic and immune toxicity^1, 16, 31, 33^; second generation nonselective mTOR inhibitors of both mTORC1 and mTORC2 have had limited clinical success perhaps due to a narrow therapeutic index^19–21^. We speculated that an mTORC1 selective inhibitor that engaged 4EBP1 would be more effective at suppressing MYC and have less nonspecific toxicity. Here we demonstrate that bi-steric selective mTORC1 inhibitors that activate 4EBP1 result in marked suppression of MYC protein levels, and drive anti-tumor activity, including regressions, in both a transgenic mouse model of MYC-driven HCC and in human PDX models of *MYC*-amplified HCC, colorectal cancer, head and neck cancer, and ovarian cancer (Fig. 6 ①). Furthermore, the present data suggest that bi-steric mTORC1-selective inhibitors are not immune suppressive or hepatotoxic, but rather, elicit robust anti-cancer immunity and cooperate with immune checkpoint therapy (Fig. 6 ②).

**Figure 6.**
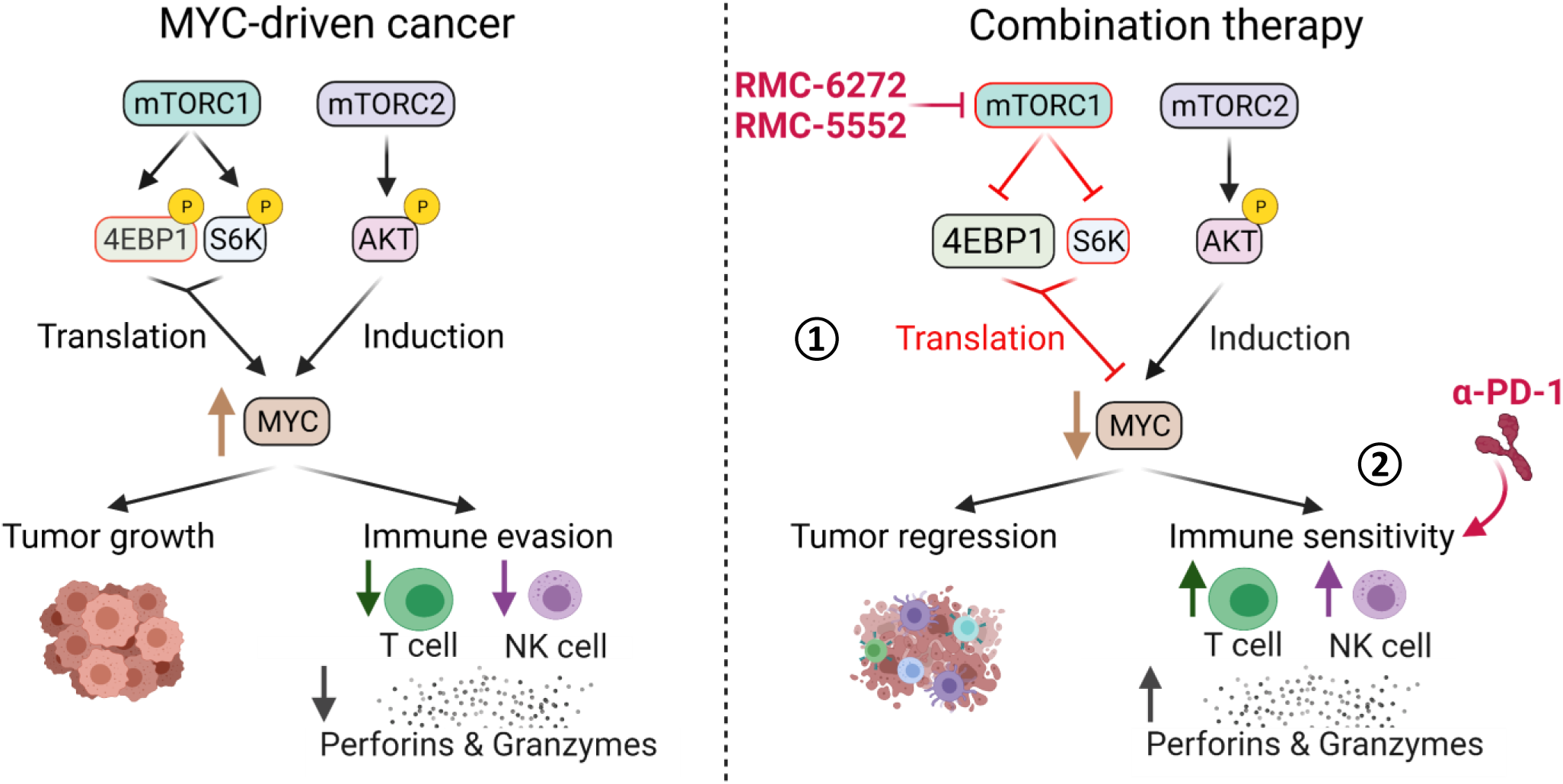
Model of anti-tumor immunity induction by the therapeutic combination of bi-steric mTORC1 inhibitors with immunotherapy. MYC-driven cancers have high p4EBP and pS6K levels which increase MYC translation inducing tumor growth and immune evasion characterized by decreased abundance of CD4+ T cells and NKp46+ NK cells; and low levels of perforins and granzyme B (left). RMC-6272 specifically inhibits the mTORC1 pathway, but not the mTORC2 pathway, resulting in reduced p4EBP1and pS6RP *in vivo*. This in turn, reduces MYC translation (right, ①) inducing tumor regression and immune activation enhancing recruitment of CD4+ T cells and NKp46+ NK cells. Addition of α-PD-1 therapy to RMC-6272-treated HCC tumors induces further immune cell activation, degranulation, and the release of perforins and granzyme B (right, ②).

mTORC1 inhibition activates 4EBP1, thereby decreasing MYC protein expression and producing an anti-tumor effect^1, 10–13^. Using representative bi-steric mTORC1-selective inhibitors to conduct preclinical studies in our MYC-driven HCC mouse model, we found that pharmacological inhibition of mTORC1 is very similar to genetic MYC inactivation. Our results suggest a model whereby mTORC1 and p4EBP1 play a key role in regulating MYC protein expression serving as a nodal point for the maintenance of a cancer phenotype (see Figs. 2-3). We also found that high *EIF4EBP1* expression and high MYC signaling are associated with poorer survival in human HCC patients when compared with patients with low *EIF4EBP1* expression and low MYC signaling (see Fig. 2b). Thus, we expanded the proof-of-principle studies with RMC-6272 and showed that RMC-5552, the bi-steric mTORC1-selective inhibitor currently undergoing clinical testing (NCT04774952), has anti-tumor activity against a panel of PDX tumors with *MYC* amplification.

Selective mTORC1 inhibition elicits immune activation and cooperates with α-PD-1 to induce tumor regression in a transgenic mouse model of MYC-driven HCC. We propose this immune activation occurs through two mechanisms:

1) Depletion of MYC *in vivo* may result in the inhibition of cell intrinsic programs that promote cell growth including cancer cell proliferation^61^ and cell cycle progression^62^, and 2) in the induction of programs that enhance cell death including apoptosis^35^ and decreased metabolism and ribosomal biosynthesis^40^. Dying cancer cells release damage associated molecular patterns (DAMPs) which promote innate anti-tumor immunity and antigen presentation by antigen-presenting cells (APCs) which prime T cells inducing their recruitment to the tumor^63, 64^. Moreover, MYC depletion *in vivo* by selective mTORC1 inhibitors may reduce the expression of immune checkpoint receptors (ICRs)^37^ on cancer cells further enhancing tumor recruitment of adaptive immune cells.

Here, we showed that inhibiting mTORC1 depletes MYC eliciting immune activation as measured by GSEA and GO Term enrichment analyses of RNA-seq data, CIBERSORT^46^ and TIMER2.0^47^, and CODEX^55, 65^. This includes the recruitment and activation of CD4+ T cells^66^, maturation and activation of NK cells^67^, and prevention of M2 macrophage polarization^8, 44^ (Fig. 4). In contrast, nonselective mTOR inhibition has been shown to cause T cell anergy^25, 68^, decrease NK cell proliferation^26^, enhance T_reg_ differentiation and expansion^28, 29^, and reduce expression of granzyme B and interferon gamma by decreasing TCR/mTORC1/IL-2R signaling^23, 24^. Second, nonselective mTOR inhibitors reduce AKT activation in both T and NK cells, which would prevent mTORC1 activation and thereby immune cell proliferation^30, 69^. Thus, bi-steric mTORC1-selective inhibitors most likely activate immune responses both by reducing MYC protein expression resulting in reduced ICR expression and DAMP release and by not inhibiting AKT, contributing to enhanced immunity.

Finally, we found that RMC-6272 cooperates with immune checkpoint inhibitors to induce tumor regression in the aforementioned mouse model of MYC-driven HCC. Earlier generations of mTOR inhibitors have been shown to be immunosuppressive^18, 22, 23–30, 31^, raising the concern that bi-steric mTOR inhibitors may interfere with immune therapies including immune checkpoint inhibitors. Current approved tyrosine kinase inhibitors for HCC, including sorafenib, have shown a limited ability to induce immune activation^70–73^ and to synergize with immune checkpoint inhibitors in HCC^74–77^. Here, we first reported that RMC-6272 alone induces immune activation in MYC-driven HCC tumors performing significantly better than sorafenib (Fig. 4). Second, we showed that RMC-6272 transformed the tumor immune microenvironment and synergized with α-PD-1 to elicit sustained tumor regression, increased T and NK cell recruitment, and the release of perforins and granzymes (Fig. 5). Selective mTORC1 inhibition via RMC-6272 most likely sensitizes MYC-driven tumors to α-PD-1 immunotherapy by overcoming MYC-driven immune evasion^8, 78^. Therefore, bi-steric mTORC1-selective inhibitors have the potential to clinically improve immune checkpoint inhibitor responses in patients with MYC-overexpressing cancers. In sum, our preclinical results support the conclusion that targeting mTORC1 has therapeutic potential in the treatment of MYC-driven cancers alone and in combination with immune checkpoint inhibition, concepts that inform the clinical testing of RMC-5552 (NCT04774952).

## Supporting information

Supplemental_Material_Mahauad-Fernandez_Yang_et_al

## Acknowledgments

We thank the Stanford Division of Oncology, and the Stanford Department of Medicine for scientific advice. We thank Delaney Sullivan for helping us running signscore and running CIBERSORT. We thank Pauline Chu in the Department of Pathology at Stanford University for helping us with tissue sectioning for immunohistochemical studies. We thank the Translational Applications Service Center (TASC) and especially Joanna Liliental and Shailja Patel for their help with Nanoimmunoassay. We thank the Stanford Cell Sciences Imaging Facility (CSIF) and Anum Khan for sample processing for CODEX. We also thank the Stanford Animal Diagnostics Laboratory and Animal Histology Service for analyses of serum samples and generation of frozen slides, respectively. We thank Revolution Medicines’ colleagues Elsa Quintana, Pete Wildes, Bianca Lee, Jingjing Jiang for key scientific input and review, as well as Ethan Ahler and Yevgeniy Gindin for RNAseq data processing. Furthermore, we thank Ensigna Biosystems for IHC support, and GenenDesign, WuXi AppTec, and Charles River Discovery Research Services for PDX studies. Figures were made using Biorender and the Qlucore software was used to analyze and generate figures from RNA-seq data.

## Authors’ contributions

WDMF, YCY, JAMS, MS, and DWF conceived the study. MS, WDMF, and YCY developed the experimental methodology. WDMF, YCY, IL, JP, LY, and JWE conducted experiments. WDMF performed experiments, analyzed data and generated figures. WDMF wrote the manuscript. JAMS, MS, GLB, AG, DWF, WDMF and YCY edited the manuscript. JAMS and DWF provided administrative, technical, and material support. JAMS, MS and DWF supervised the study. All authors read and approved the final manuscript.

## Supplemental Figure Legends

**Supplemental Figure 1. Low MYC signaling and EIF4EBP1 levels in human HCC tumors are predictive of worse survival. a-b**, Kaplan Meier survival plots of **a**, progression-free survival and **b**, disease-free survival of human HCC patients from TCGA LIHC after stratifying them into those expressing high and low MYC signature (MYC^Sig^), using the Hallmarks of MYC targets geneset and signscore, and expressing high or low *EIF4EBP1* levels. Log rank (Mantel-Cox) test was used for Kaplan-Meier survival analyses to compare indicated groups. Significance was taken at P<0.05 (*) and P<0.005 (***). ns = not significant

**Supplemental Figure 2. The bi-steric mTORC1-selective inhibitor RMC-6272 potently reduces MYC and p4EBP1 levels. a**, Relative protein levels, determined by nanoimmunoassay (NIA), and **b**, determined by immunohistochemistry (IHC) of hMYC, p4EBP1, pS6RP, and pAKT; and, **c**, representative NIA western blot-like images and **d**, quantification of ERK1 and 2 and their different phosphorylated isoforms from HCC tumors in Lap-tTA/Tet-O-MYC mice treated for 7 days with a vehicle control (5/5/90 Transcutol/Solutol HS 15/Water) injected intraperitoneally once weekly, doxycycline at 100 µg/ml in water (MYC Off), sorafenib at 30 mg/kg/day given via oral gavage, or two bi-steric mTORC1-selective inhibitors RMC-4627 and RMC-6272 at 10 mg/kg injected intraperitoneally once weekly. At day 7, mice were treated with one more dose of their respective treatment and euthanize 24 hours after. Error bars represent minimum and maximum whiskers for panels **a** and **b** and SEM for panel **d**. Significance was taken at P<0.05 (*), P<0.01 (**), P<0.005 (***), and P<0.001 (****). ns = not significant.

**Supplemental Figure 3. RMC-6272 reduces the expression of mTORC1- and MYC-target genes without affecting expression of genes regulated by mTORC2 or MAPK. a**, Heat maps of genes and **b**, GSEA plots of the following genesets: hallmarks of MYC targets, hallmarks of mTORC1 signaling, hallmarks of PI3K, AKT, mTOR signaling, and activated MAPK activity from RNA-seq data of HCC tumors in Lap-tTA/Tet-O-MYC mice treated for 7 days with a vehicle control (5/5/90 Transcutol/Solutol HS 15/Water), doxycycline at 100 µg/ml in water (MYC Off), Sorafenib at 30 mg/kg/day given via oral gavage or the selective mTORC1 inhibitors RMC-4627 or RMC-6272 given intraperitoneally at 10 mg/kg once weekly. At day 7, mice were treated with one more dose of their respective treatment and euthanize 24 hours after.

**Supplemental Figure 4. Effect of RMC-6272 on the expression of genes regulated by oncogenic signaling pathways common to HCC. a-h**, Heat maps (top) and GSEA plots (bottom) of the following genesets: **a**, hallmarks of MYC targets, **b**, canonical WNT signaling pathway, **c**, neoplastic transformation by CCND1 (WNT) and MYC, **d**, RAS signaling pathway, **e**, HRAS oncogenic signature, **f**, KRAS signaling downregulated genes, **g**, reactome signaling by BRAF and RAF fusions, and **h**, PI3K and AKT signaling pathway from RNA-seq data of HCC tumors in Lap-tTA/Tet-O-MYC mice treated for 7 days with a vehicle control (5/5/90 Transcutol/Solutol HS 15/Water), doxycycline at 100 µg/ml in water (MYC Off), Sorafenib at 30 mg/kg/day given via oral gavage or the selective mTORC1 inhibitors RMC-4627 or RMC-6272 given intraperitoneally at 10 mg/kg once weekly. At day 7, mice were treated with one more dose of their respective treatment and euthanize 24 hours after. ES = enrichment scores.

**Supplemental Figure 5. RMC-6272 induces a durable decrease of p4EBP1, pS6RP, and MYC protein levels compared to RMC-4627. a-d**, Quantification of **a**, hMYC, **b**, p4EBP1, and **c**, pS6RP protein levels as detected by immunohistochemistry. Lap-tTA/Tet-O-MYC mice were treated for 7 days with a vehicle control (5/5/90 Transcutol/Solutol HS 15/Water) injected intraperitoneally once weekly, doxycycline at 100 µg/ml in water (MYC Off), sorafenib at 30 mg/kg/day given via oral gavage, or two bi-steric mTORC-selective inhibitors RMC-4627 and RMC-6272 at 10 mg/kg injected intraperitoneally once weekly. Mice were euthanized for tumor collection at the following time points after a final dose on Day 7: 4, 12, 24, 47, and 72 hours. Error bars represent SEM. Significance was taken at P<0.01 (**), and P<0.005 (***). ns = not significant.

**Supplemental Figure 6. RMC-5552 significantly reduces tumor growth in PDX models of several human cancer types.** Tumor volume over time of a **a**, colorectal cancer PDX, **b**, head and neck cancer PDX, and **c**, an ovarian cancer PDX with high MYC amplifications treated with a vehicle control (5/5/90 Transcutol/Solutol HS 15/Water) or RMC-5552 given intraperitoneally at 3 or 10 mg/kg/week for 28 days. Tumor volumes at the end of study were compared between control and treatment groups by two-way ANOVA Dunnet’s multiple comparison test. Significance was taken at P<0.05 (*), P<0.01 (**), and P<0.005 (***), ns = not significant.

**Supplemental Figure 7. RMC-6272 induces regression of high burden MYC-driven tumors.** Representative **a**, MRI images, **b**, 3D HCC tumors, and **c-g**, tumor volume over time of HCC tumors in Lap-tTA/Tet-O-MYC mice before (Day 0) and after (Day 7) treatment with **c**, a vehicle control (5/5/90 Transcutol/Solutol HS 15/Water) injected intraperitoneally once weekly, **d**, doxycycline at 100 µg/ml in water (MYC Off), two pre-clinical bi-steric selective mTORC1 inhibitors **e**, RMC-4627 and **f**, RMC-6272 given intraperitoneally at 10 mg/kg/week, and **g**, sorafenib given via oral gavage at 30 mg/kg/day.

**Supplemental Figure 8. Bi-steric mTORC1 inhibitors do not induce body weight loss or liver toxicity in MYC-driven HCC model. a**, body weight change at Day 7 depicted as a percent over initial weight at Day 0 (before treatment), and **b**, levels of circulating liver enzymes alanine aminotransferase (ALT), aspartate transaminase (AST), alkaline phosphatase (Alk. Phos.), bilirubin, gamma-glutamyl transferase (GGT); and, **c**, cholesterol, **d**, triglycerides, **e**, sodium, **f**, high-density lipoprotein (HDL), **g**, low-density lipoprotein (LDL), **h**, chloride, **i**, glucose, **j**, albumin, **k**, creatinine, **l**, blood urea nitrogen (BUN), **m**, calcium, **n**, globulin, **o**, potassium **p**, CO_2_, and **q**, total protein levels from serum of Lap-tTA/Tet-O-MYC mice treated with a vehicle control (5/5/90 Transcutol/Solutol HS 15/Water) injected intraperitoneally once weekly, doxycycline, sorafenib 30 mg/kg daily orally, once weekly intraperitoneal injection of RMC-4627, or RMC-6272 for 7 days. Gray squares depict the range of normal values for specific factors in wild-type mice. Error bars represent minimum and maximum whiskers. Significance was taken at P<0.05 (*), P<0.01 (**), P<0.005 (***), and P<0.001 (****). ns = not significant.

**Supplemental Figure 9. Bi-steric mTORC1 inhibitors are not associated with hyposplenism in MYC-driven HCC model.** Spleen sizes of Lap-tTA/Tet-O-MYC mice treated for 7 days with a vehicle control (5/5/90 Transcutol/Solutol HS 15/Water) injected intraperitoneally once weekly, RMC-4627 or RMC-6272 at 10 mg/kg injected intraperitoneally once weekly, or sorafenib at 30 mg/kg/daily via oral gavage. Error bars represent SEM. Significance was taken at P<0.05 (*).

**Supplemental Figure 10. MYC signature and *EIF4EBP1* downregulation is predictive of immune reactivation.** Heat map depicting relative abundance of different immune populations in MYC^Sig^ and *EIF4EBP1* high *versus* low HCC tumors (TCGA LIHC) using CIBERSORT.

**Supplemental Figure 11. RMC-6272 in combination with α-PD-1 drives tumor regression in a MYC-driven HCC mouse model. a**, Representative 2D MRI images over time, **b**, representative 3D HCC tumors over time, **c-f**, tumor volume over time, and **g**, gross images of HCC tumors in Lap-tTA/Tet-O-MYC mice before treatment (Day 0) and after weekly treatments for 28 days with **c**, a vehicle control (5/5/90 Transcutol/Solutol HS 15/Water) injected intraperitoneally once weekly, **d**, α-PD-1 injected intraperitoneally at 300 µg triweekly, **e**, RMC-6272 injected intraperitoneally at 10 mg/kg/week, or **f**, a combination of RMC-6272 with α-PD-1. Scale bar in panel **g** is 10 mm.

**Supplemental Figure 12. Effects of RMC-6272 in combination with α-PD-1 in levels of several circulating factors. a**, Levels of circulating liver enzymes: alanine aminotransferase (ALT), aspartate transaminase (AST), alkaline phosphatase (Alk. Phos.), bilirubin, and gamma-glutamyl transferase (GGT) from Lap-tTA/Tet-O-MYC mice treated with vehicle, RMC-6272, α-PD-1, or RMC-6272 with α-PD-1 for 28 days. **b**, Body weight change over time depicted as a percent over initial weight at Day 0 (before treatment). **c**-**q**, Levels of circulating **c**, cholesterol, **d**, triglycerides, **e**, sodium, **f**, potassium, **g**, high-density lipoprotein (HDL), **h**, low-density lipoprotein (LDL), **i**, chloride, **j**, CO_2_, **k**, glucose, **l**, albumin, **m**, creatinine, **n**, total protein, **o**, blood urea nitrogen (BUN), **p**, calcium, and **q**, globulin levels from serum of Lap-tTA/Tet-O-MYC mice before treatment (Day 0) and after weekly treatments for 28 days with a vehicle control (5/5/90 Transcutol/Solutol HS 15/Water) injected intraperitoneally once weekly, α-PD-1 injected intraperitoneally at 300 µg triweekly, RMC-6272 injected intraperitoneally at 10 mg/kg/week, or a combination of RMC-6272 with α-PD-1. Gray squares depict the range of normal values for specific factors in wild-type mice. Error bars represent minimum and maximum whiskers. Significance was taken at P<0.05 (*) and P<0.01 (**). ns = not significant.

