## Supplemental_Material_Mahauad-Fernandez_Yang_et_al for "Bi-steric mTORC1-selective Inhibitors activate 4EBP1 reversing MYC-induced tumorigenesis and synergize with immunotherapy"

### SUPPLEMENTAL FIGURE 1

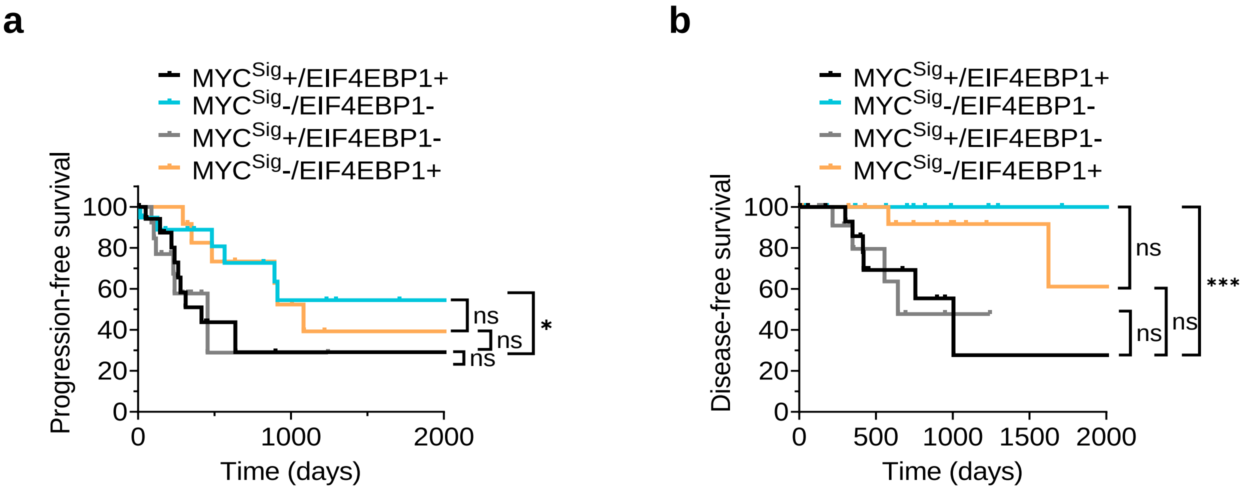

### SUPPLEMENTAL FIGURE 2

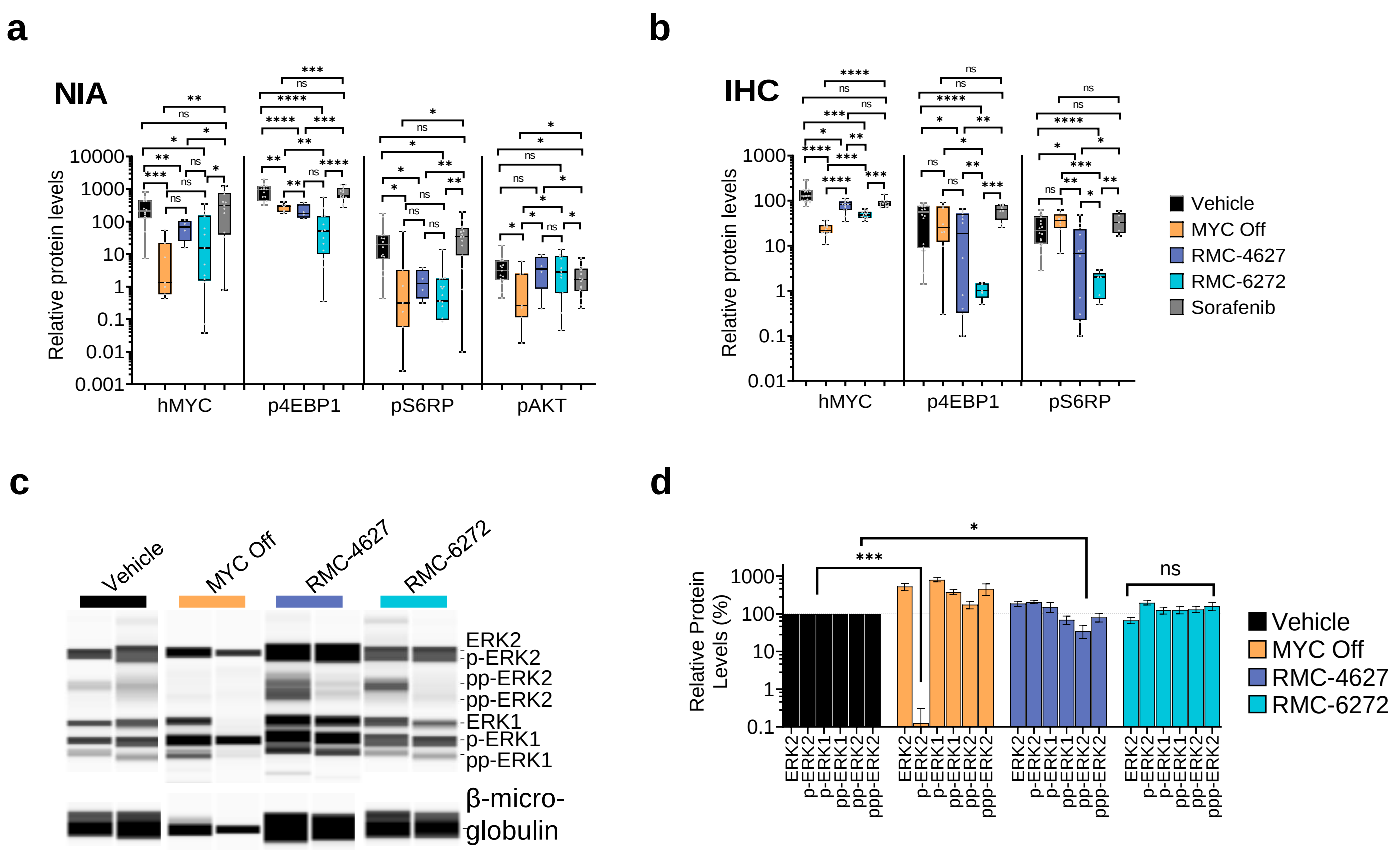

### SUPPLEMENTAL FIGURE 3

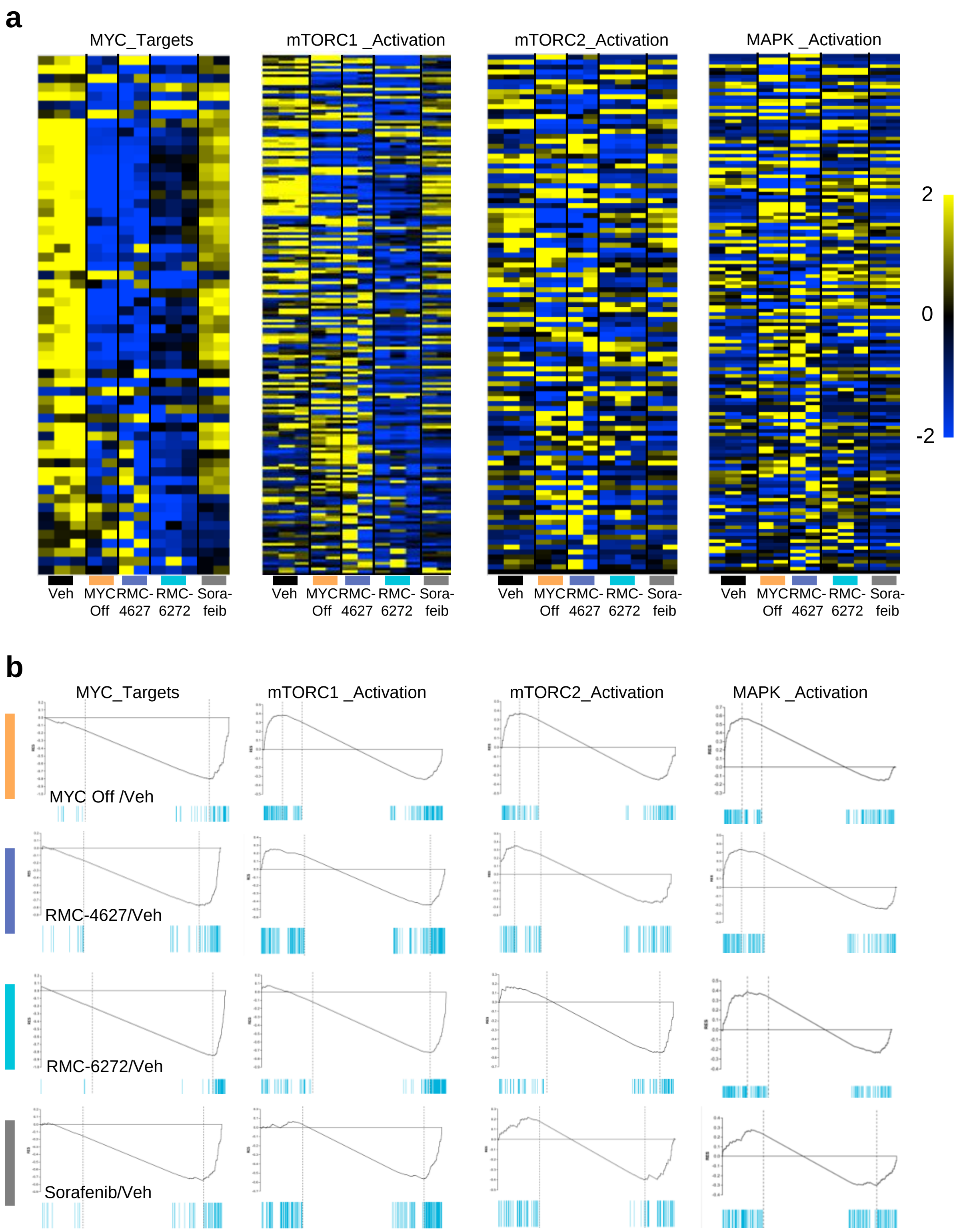

### SUPPLEMENTAL FIGURE 4

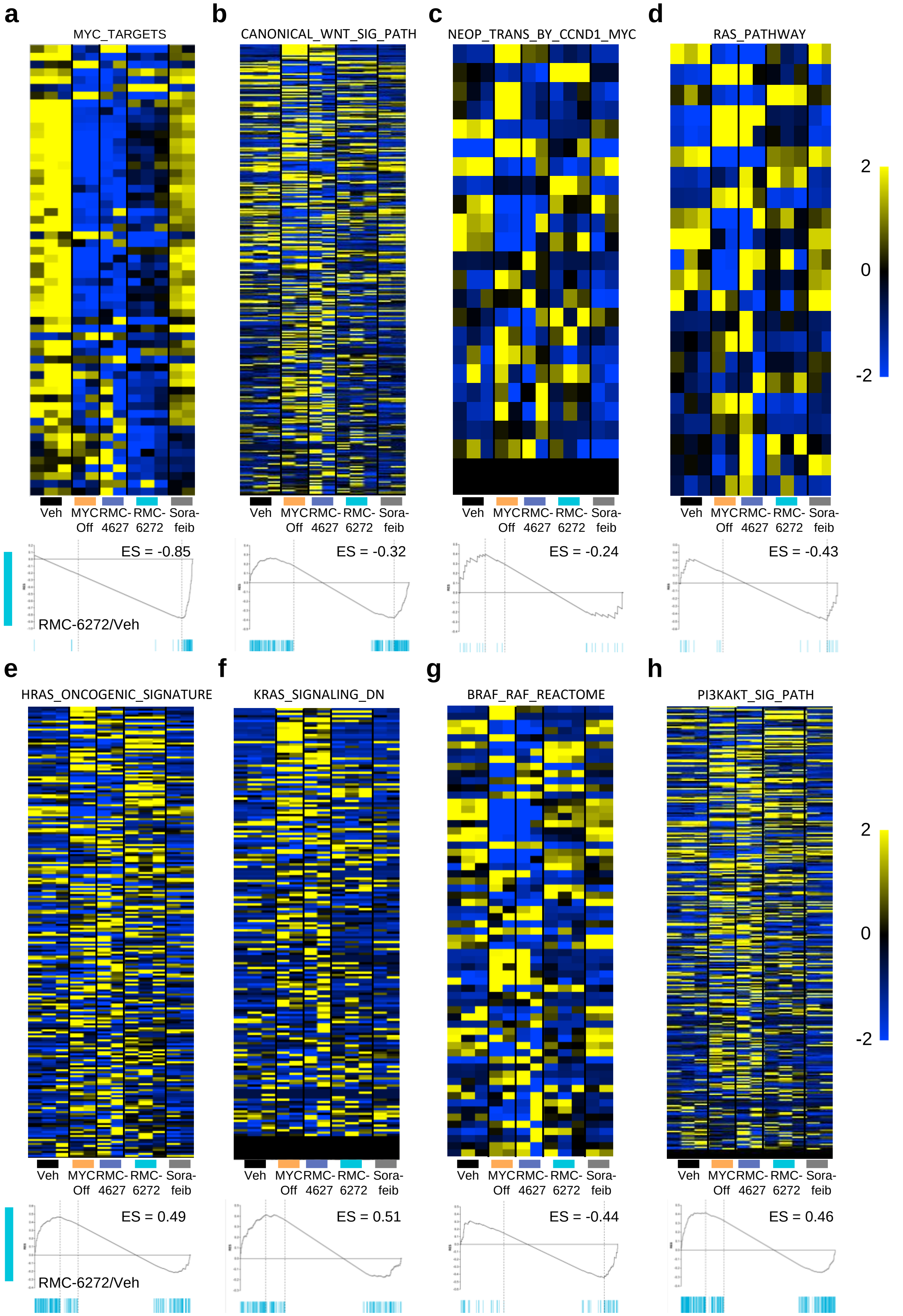

### SUPPLEMENTAL FIGURE 5

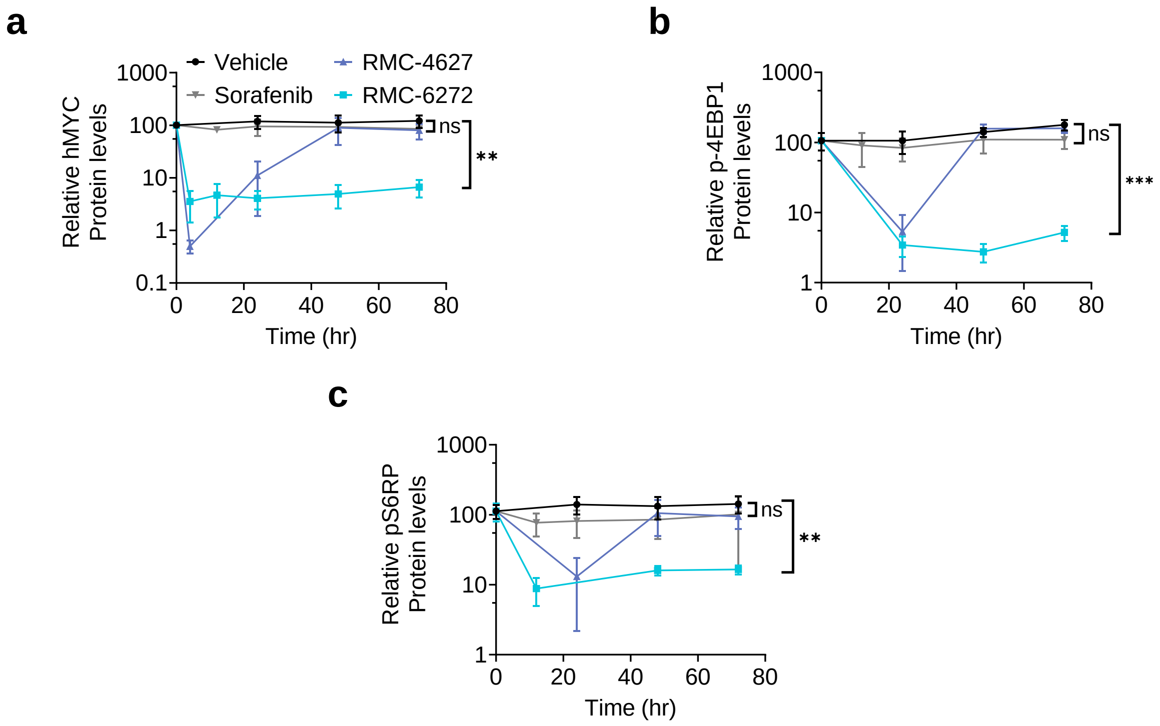

### SUPPLEMENTAL FIGURE 6

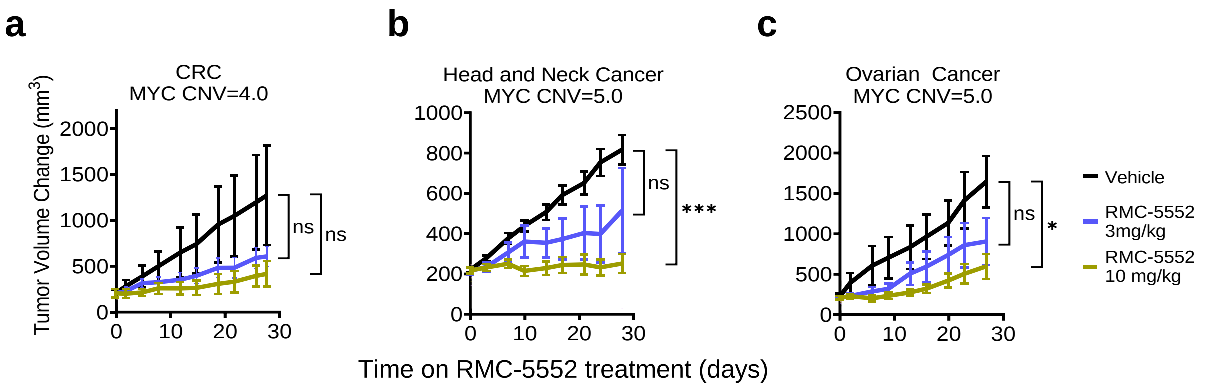

### SUPPLEMENTAL FIGURE 7

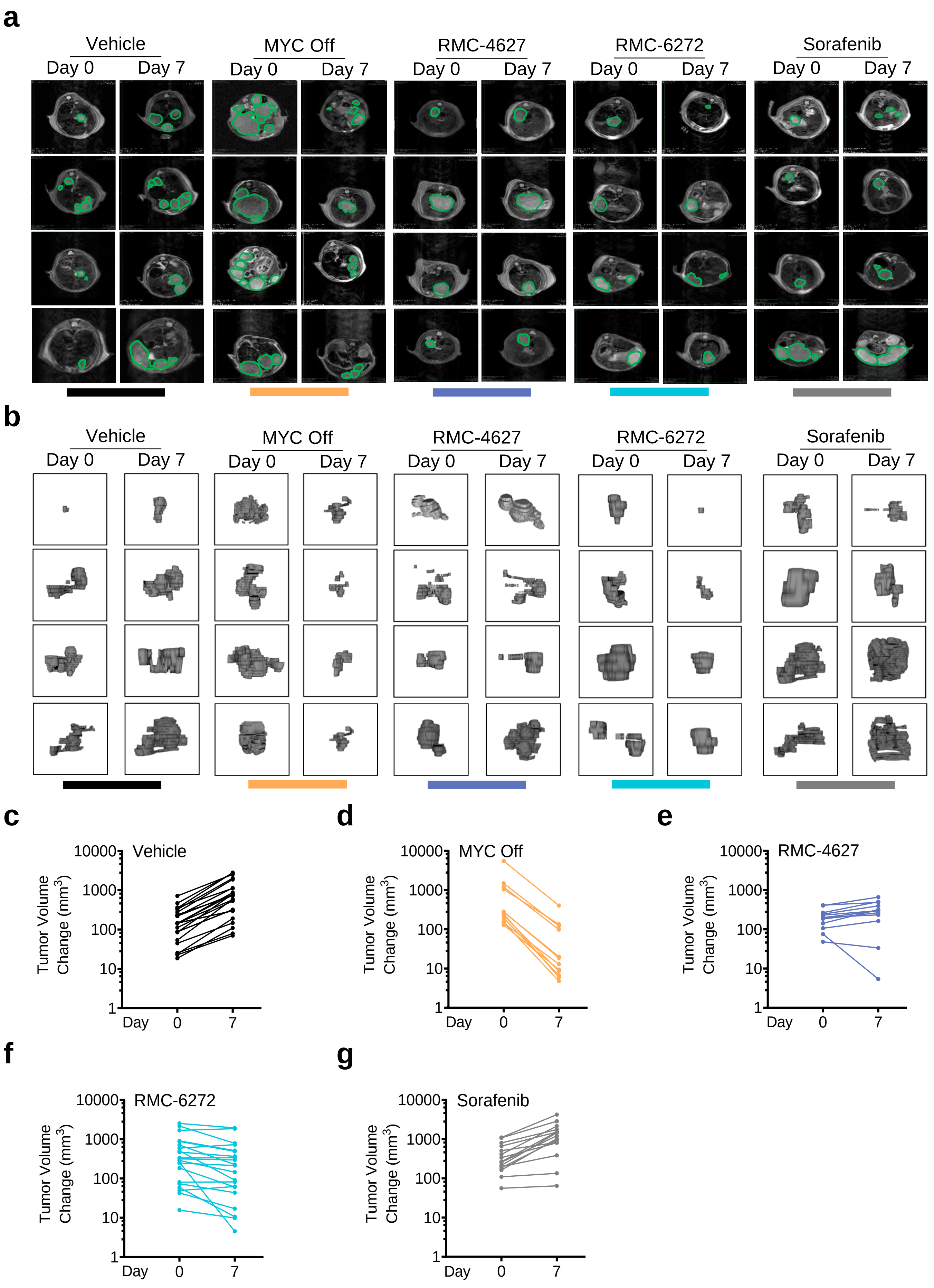

### SUPPLEMENTAL FIGURE 8

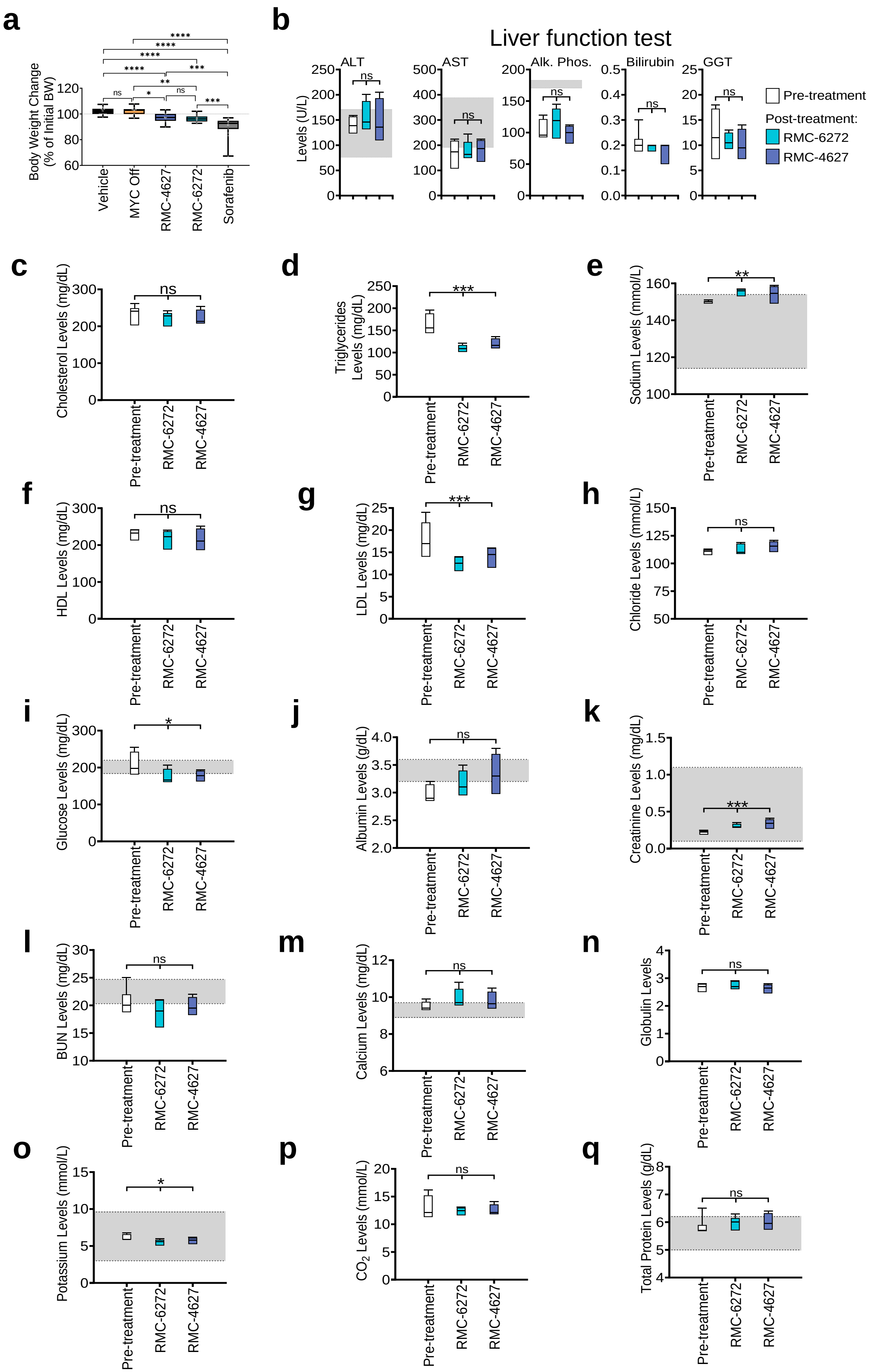

### SUPPLEMENTAL FIGURE 9

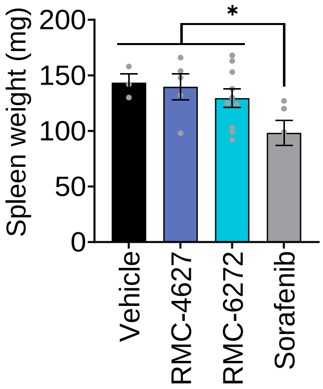

### SUPPLEMENTAL FIGURE 10

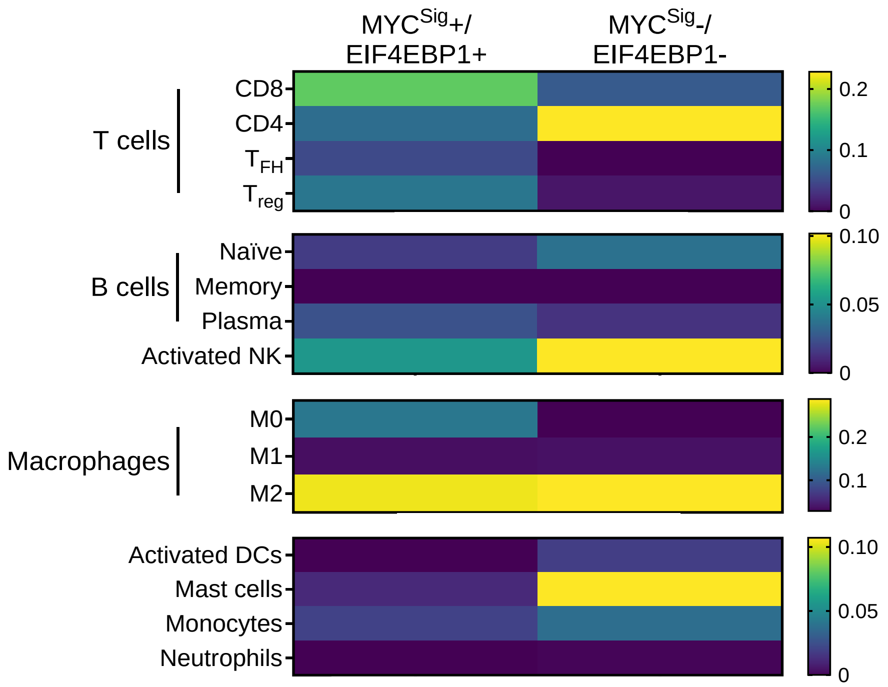

### SUPPLEMENTAL FIGURE 11

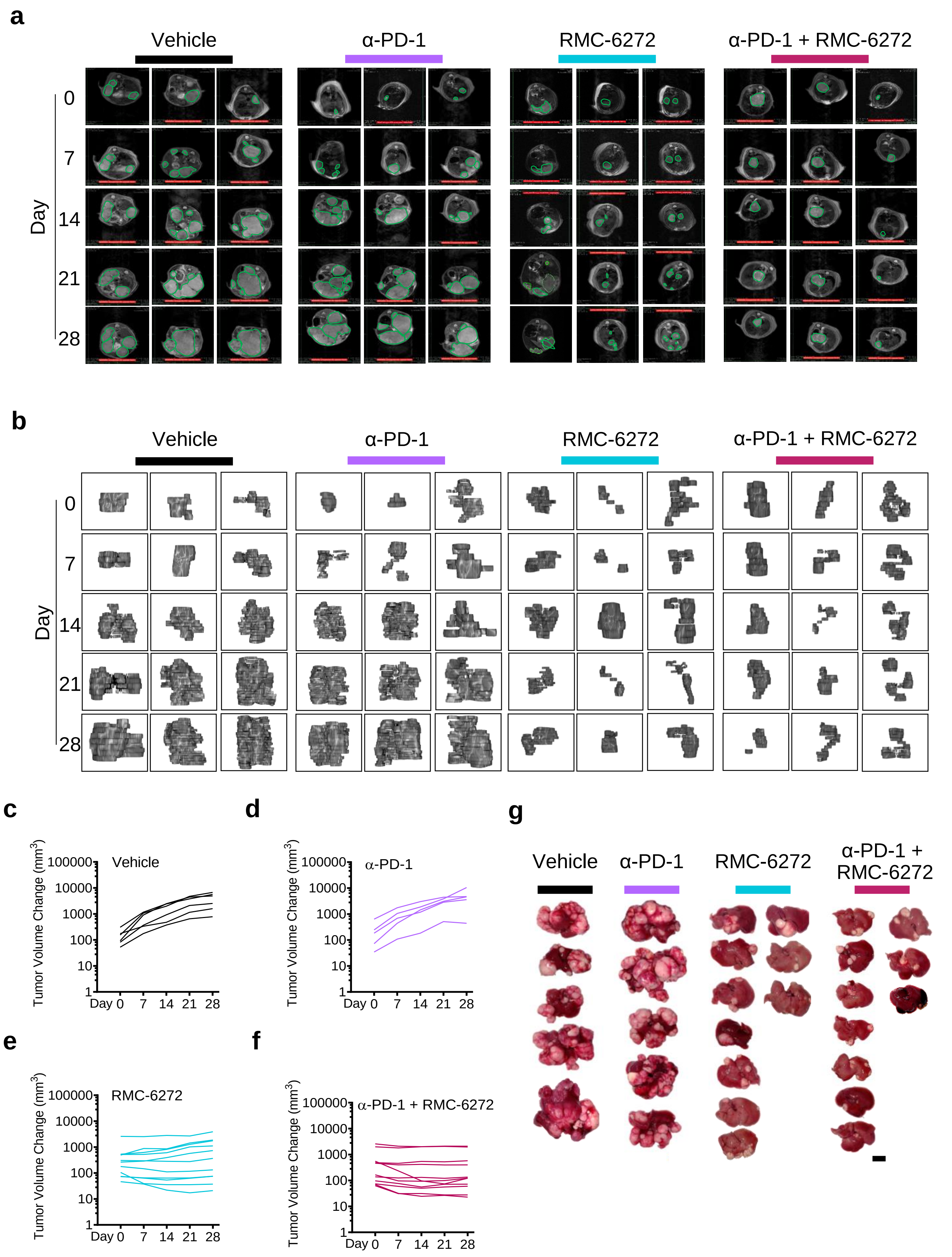

### SUPPLEMENTAL FIGURE 12

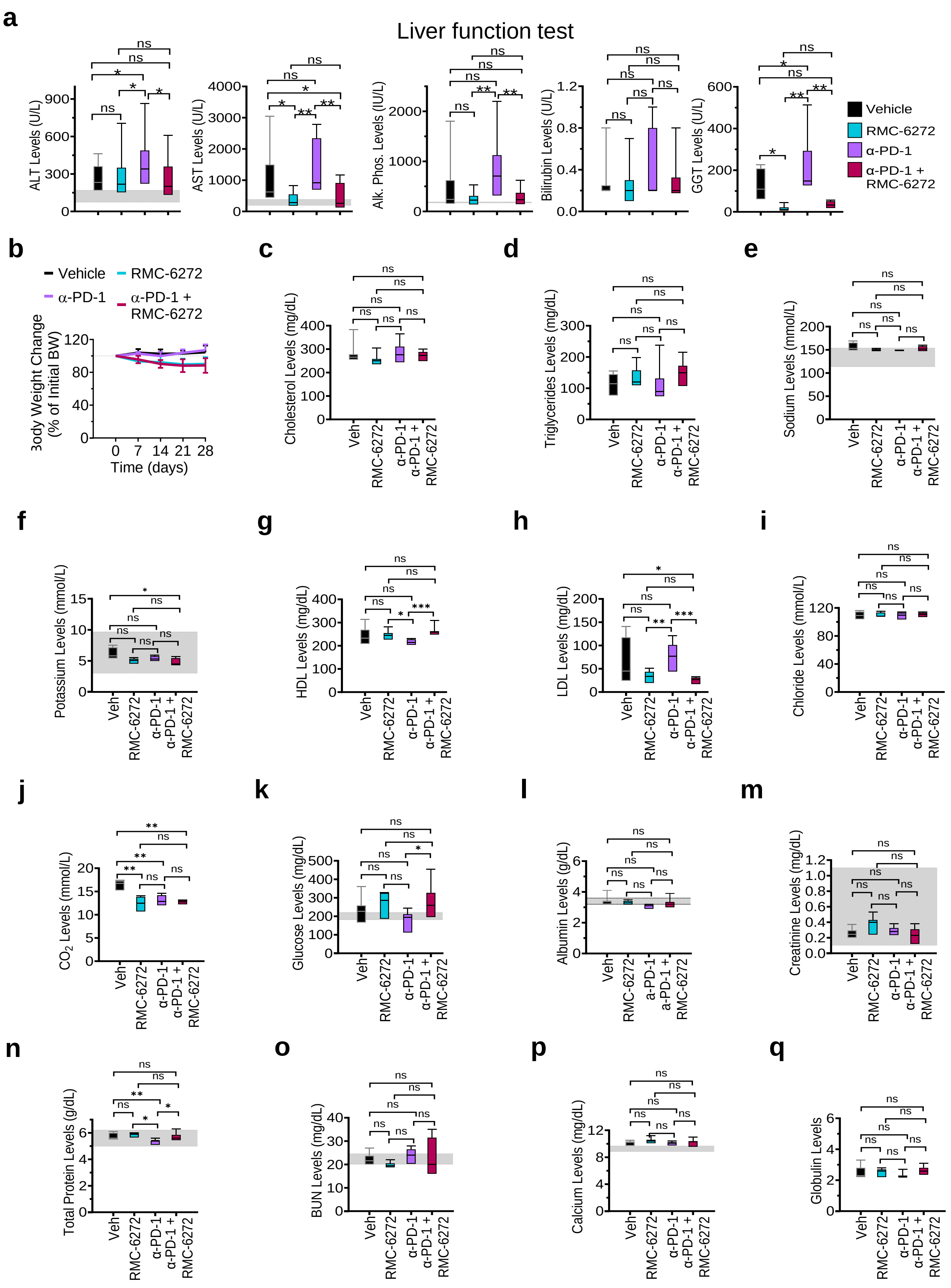

**Supplemental Table 1. Average tumor volumes of HCC mice pre and post 7 day treatment**

| Treatment group | Average tumor volume (mm^3) | | p-value | | Number of mice/group |
|---|---|---|---|---|---|
| | Day 0 | Day 7 | Day 0 | Day 7 | |
| Vehicle | 187.55 ± 163.11 | 896.89 ± 837.60 | N/A | 0.003452455 | n=25 |
| MYC OFF (Doxycyline) | 906.72 ± 1525.79 | 70.40 ± 115.54 | 0.131432467 | 5.21541E-05 | n=12 |
| RMC-4627 | 208.89 ± 114.14 | 304.85 ± 189.93 | 0.648620268 | 0.002223853 | n=12 |
| RMC-6272 | 588.68 ± 706.71 | 393.56 ± 550.16 | 0.01854044 | 0.00016512 | n=21 |
| Sorafenib | 423.03 ± 339.253 | 1390.79 ± 1075.17 | 0.021460095 | 0.140661694 | n=15 |

| Treatment group | Average % change | | p-value | | Number of mice/group |
|---|---|---|---|---|---|
| | Day 0 | Day 7 | Day 0 | Day 7 | |
| Vehicle | 100 ± 0 | 495.42 ± 232.08 | N/A | 1.51976E-06 | n=25 |
| MYC OFF (Doxycyline) | 100 ± 0 | 6.63 ± 3.01 | N/A | 7.63872E-08 | n=12 |
| RMC-4627 | 100 ± 0 | 135.78 ± 55.99 | N/A | 2.38912E-05 | n=12 |
| RMC-6272 | 100 ± 0 | 64.86 ± 35.90 | N/A | 5.47811E-07 | n=21 |
| Sorafenib | 100 ± 0 | 354.95 ± 194.43 | N/A | 0.07257789 | n=15 |

p-values are relative to Vehicle Day 0. p-value calculated using a t-test with two-tailed distributions and two samples of unequal variances

**Supplemental Table 2. Weekly average tumor volumes of HCC mice over a 4 week treatment period**

| Treatment group | Average tumor volume (mm^3) | | | | | p-value | | Number of mice/group |
|---|---|---|---|---|---|---|---|---|
| | Day 0 | Day 7 | Day 14 | Day 21 | Day 28 | Day 0 | Day 28 | |
| Vehicle | 143.70 ± 91.97 | 615.19 ± 348.13 | 1331.81 ± 806.30 | 2475.85 ± 1152.31 | 2975.14 ± 1833.53 | N/A | 0.01364698 | n=6 |
| anti-PD-1 | 234.01 ± 239.80 | 598.24 ± 402.90 | 1426.98 ± 1049.95 | 2708.07 ± 1399.66 | 3176.67 ± 300.34 | 0.462875866 | 0.841509126 | n=5 |
| RMC-6272 | 506.63 ± 715.89 | 445.65 ± 578.40 | 512.94 ± 440.92 | 668.59 ± 486.23 | 844.76 ± 300.28 | 0.33484555 | 0.0077166 | n=11 |
| anti-PD-1 + RMC-6272 | 606.21 ± 858.78 | 410.10 ± 872.27 | 419.61 ± 690.36 | 417.7 ± 660.36 | 446.89 ± 650.15 | 0.106343056 | 0.006372064 | n=11 |

| Treatment group | Average % change | | | | | p-value | | Number of mice/group |
|---|---|---|---|---|---|---|---|---|
| | Day 0 | Day 7 | Day 14 | Day 21 | Day 28 | Day 0 | Day 28 | |
| Vehicle | 100 ± 0 | 429.93 ± 345.75 | 922.85 ± 822.38 | 1731.27 ± 1535.40 | 2080.39 ± 1524.02 | N/A | 0.074781164 | n=6 |
| anti-PD-1 | 100 ± 0 | 259.65 ± 164.12 | 620.34 ± 628.05 | 1258.48 ± 1373.65 | 1367.873 ± 2017.24 | N/A | 0.27906449 | n=5 |
| RMC-6272 | 100 ± 0 | 87.89 ± 32.46 | 101.05 ± 43.50 | 131.97 ± 84.56 | 166.76 ± 113.30 | N/A | 0.007399109 | n=11 |
| anti-PD-1 + RMC-6272 | 100 ± 0 | 67.65 ± 17.115 | 69.22 ± 29.970 | 68.91 ± 42.203 | 73.72 ± 35.194 | N/A | 0.006304996 | n=11 |

p-values are relative to Vehicle Day 0. p-value calculated using a t-test with two-tailed distributions and two samples of unequal variances
